# Sex-dependent effects of a high-fat diet on the hypothalamic response in mouse

**DOI:** 10.1101/2024.09.10.612192

**Authors:** Virginie Dreux, Candice Lefebvre, Charles-Edward Breemeersch, Colin Salaün, Christine Bôle-Feysot, Charlène Guérin, Pierre Déchelotte, Alexis Goichon, Moïse Coëffier, Ludovic Langlois

**Author notes:** Both authors contributed equally (Moïse Coëffier, Ludovic Langlois). **Corresponding author:** Dr Ludovic Langlois, Inserm UMR1073 – UFR santé – 22 boulevard Gambetta – F-76000 Rouen, France –.

## Abstract

Sex differences in rodent models of diet-induced obesity are still poorly documented, particularly regarding how central mechanisms vary between sexes in response to an obesogenic diet. Here, we wanted to determine whether obesity phenotype and hypothalamic response differed between male and female C57Bl/6J mice when exposed to a high-fat diet (HFD). Mice were exposed either a free 60% HFD or standard diet first for both a long-(14 weeks) and shorter-periods of time (3, 7, 14 and 28 days). Analysis of the expression profile of key neuronal, glial and inflammatory hypothalamic markers was performed using RT-qPCR. In addition, astrocytic and microglial morphology was examined in the arcuate nucleus. Monitoring of body weight and composition revealed that body weight and fat mass gain appeared earlier and was more pronounced in male mice. After 14 weeks of HFD exposure, normalized increase of body weight reached similar levels between male and female mice. Overall, both sexes under HFD displayed a decrease of orexigenic neuropeptides expression and an increase in POMC gene expression was observed only in female mice. In addition, changes in the expression of hypothalamic inflammatory markers were relatively modest. We also reported that the glial cell markers expression and morphology were affected by HFD in a sex- and time dependent manner, suggesting a more pronounced glial cell activation in female mice. Taken together, these data show that male and female mice responded differently to HFD exposure, both on short- and long-term and suggest that a strong inflammatory hypothalamic profile is not systematically present in DIO models. Nevertheless, in addition to these present data, the underlying mechanisms should be deciphered in further investigations.

## 1. Background

The World Health Organization reported that the prevalence of obesity has been rising worldwide for several decades reaching 13% of adults (11% of men and 15% of women) in 2016. Obesity is associated with many comorbidities (cardiovascular diseases, metabolic syndrome, anxio-depressive disorders), thus representing a major public health problem (Tutor et al., 2023). This multifactorial disease is characterized by an energy imbalance mostly due to changes in the dietary and physical activity patterns of the modern population. In particular, the consumption of energy-dense foods and excess lipids leads to low-grade chronic inflammation, the principal component of obesity (Gregor and Hotamisligil, 2011).

In periphery, this inflammatory response is orchestrated by metabolic organs (including adipose tissue, liver, muscle and pancreas) and involves several pathways (Gregor and Hotamisligil, 2011). One of them is related to the expansion and hypertrophy of adipocytes, which release proinflammatory cytokines and free fatty acids in systemic circulation, activating the immune system (Redinger, 2007). The role of gut microbiota is also discussed *via* microbiota-gut-brain communication (Solas et al., 2017). The inflammatory state disrupts especially the hypothalamic function (De Souza et al., 2005; Thaler et al., 2012).

Given the critical role of the hypothalamus in appetite control and energy expenditure (Luquet, 2008; Schwartz et al., 2000), extensive studies have been carried out to examine the link between hypothalamic inflammation and the pathophysiology of obesity and associated metabolic disorders (Bhusal et al., 2022; Cai and Liu, 2011; Jais and Brüning, 2017; Thaler and Schwartz, 2010). These studies demonstrated that the diet-related brain inflammatory response precedes weight gain or obesity development (Thaler et al., 2012). Indeed, hypothalamic inflammatory response can be detected at the postprandial time scale (Cansell et al., 2021). Although the underlying biological processes and mediators involved still remain unclear, glial cells such as microglia and astrocytes surrounding neurons located in brain areas linked to food intake regulation have been proposed as major cellular contributors to hypothalamic inflammation (Lee et al., 2020; Rahman et al., 2018). Especially, a reactive gliosis was described in response to the consumption of HFD. This glial signature is characterized by morphological and phenotypic changes (increased ionized calcium-binding adapter molecule 1, Iba1 and glial fibrillary acidic protein, GFAP expressions) associated with functional dysregulation (Lee et al., 2020; Rahman et al., 2018; Sewaybricker et al., 2022; Valdearcos et al., 2014).

Despite the female predominance in obesity, most of preclinical studies have been conducted in male rodents (Baufeld et al., 2016; Leyh et al., 2021; Pistell et al., 2010). Nevertheless, some data reported sex differences, with females exhibiting an overall resistance to the obesogenic effects of HFD (Huang et al., 2020; Maric et al., 2022; Lefebvre et al., 2024). In addition, research has been focused on the effects of either long-term or short-term exposure to HFD, and often in other brain structures such as the hippocampus (de Paula et al., 2021), the ventral tegmental area (Mizoguchi et al., 2021) or the nucleus accumbens (Décarie-Spain et al., 2018). Consequently, sex differences in the hypothalamic response after different time exposure to HFD remain poorly documented within the same study.

In the present study, we aimed to evaluate how sex and the duration of exposure to HFD can influence the characteristics of the diet-induced obesity model (DIO). We investigated the hypothalamic response after long- and short-term exposure to HFD in C57BL/6J male and female mice, focusing on the assessment of expression levels of several inflammatory markers and glial cell-specific markers.

## 2. Materials and methods

### 2.1 Animal experimentation

Six-week-old C57Bl/6J male and female mice were obtained from Janvier Labs (Le Genest-Saint-Isle, France, n=24/sex, experiment 1; n=72/sex, experiment 2). Mice were socially housed (n=4/cage) in a controlled environment (20±2°C with a 12-hour light/dark cycle) with free access to food and water. After 1 week of acclimatization, male and female mice were randomized into different groups receiving for 14 weeks (experiment 1, w14) or for 3, 7, 14, and 28 days (experiment 2, d3, d7, d14, d28) either a standard diet providing 14% from fat, 27% from proteins and 59% from carbohydrates (SD, 3.34 kcal/g, 1314 formula, Altromin, Lage, Germany) or high fat diet (HFD) providing 60% kcal as fat, 20% kcal as carbohydrates and 20% kcal as protein (5.24 kcal/g, D12492i, Research Diet, New Brunswick, NJ, US, **Supplementary Table S1**). Body weight was monitored weekly, and body composition was assessed by EchoMRI (EchoMRI, Houston, TX, US) at the end of each time of diet exposure. All animal experiments were carried out in accordance with ARRIVE guidelines and the EU Directive 2010/63/EU for animal experiments. The protocols used were approved by the regional ethics committee and authorized by the French Ministry of Higher Education, Research and Innovation (authorization on APAFIS #29283-2021012114574889 v5).

### 2.2 Tissue sampling

Prior to tissue collection, mice were deeply anesthetized by intraperitoneal injection of a ketamine/xylazine solution (100 and 10 mg/kg, respectively). Blood samples were collected, centrifuged (3000*g,* 4°C for 15 min) in heparinized tubes and the plasma was frozen at -80°C. Mice were decapitated and the brains were removed on ice either for immunohistochemistry analyses or for hypothalamus dissection. The brain and hypothalamus were immediately placed in a 4% paraformaldehyde (PFA) buffered solution or frozen in liquid nitrogen and stored at - 80°C, respectively.

### 2.3 Brain slice preparation and immunofluorescence

The whole brains (n=16 for experiment 1; n=72 for experiment 2) were post-fixed for 24 h in 4% PFA buffered solution at room temperature (RT), then cryoprotected into a 30% sucrose solution (in 0.1 M phosphate-buffered saline (PBS)). Following at least 24 h of cryoprotection, the brains were frozen between -20 and -30°C in isopentane before being stored at -80°C. Serial coronal sections (20-µm-thick), located -1.22 to -2.54 mm from bregma based on brain atlas coordinates (The Mouse Brain in Stereotaxic Coordinates, 3^rd^ Edition, Franklin and Paxinos, 2008), were cut using a CM 1950 cryostat (Leica Biosystems, Ruel-Malmaison, France). Sections were mounted on Superfrost Plus microscope slides (Thermo Fisher Scientific, Portmouth, US) and then stored at -20°C.

Before immunohistochemistry, the slides were rehydrated in PBS (15 min). Then, heat-induced epitope retrieval was performed using 10 mM sodium citrate (pH 8.5) buffer for 20 min at 80°C. Next, brain slices were blocked in 5% bovine serum albumin (BSA) (Eurobio Abcys, Courtaboeuf, France) in PBS for 30 min at RT and incubated overnight at 4°C with the following antibodies: rat GFAP monoclonal immunoglobulin G (IgG, Cat. No 13-0300, 1:500, Invitrogen, Rockford, US), and rabbit IBA1 polyclonal IgG (Cat. No 019-19741, 1:2000, FUJIFILM Wako Pure Chemical, Osaka, Japan) in PBS, 0.3% Triton X-100, 0.01% NaN_3_. The next day, after being washed in PBS (3x10 min) and TNT solution (1.2% Tris; 0.9% NaCl; 1 mL Tween 20; distilled water qsp 2 L) (30 min), the slides were blocked with TNB solution for 30 min (1.2% Tris; 0.9% NaCl; 0.01% skim milk in distilled water) and incubated with Alexa Fluor 488-conjugated anti-rat IgG secondary antibody (Cat. No A11006, 1:400, Invitrogen,) and Alexa Fluor 555-conjugated anti-rabbit IgG secondary antibody (Cat. No A21428,1:400; Life Technologies,) protected from light for 2 h. After washing in TNT (3x10 min), the slides sections were mounted between the slide and coverslip with a solution of Fluoroshield with DAPI (Sigma Aldrich, Saint Louis, MO, US) and then stored at 4°C in the dark until microscopic observation. The coverslips were sealed with nail polish to prevent desiccation and movement of the samples under the microscope. Controls without primary antibodies were used to check the absence of nonspecific coupling of secondary antibodies.

### 2.4 Image acquisition

#### 2.4.1 Cell counting

For the experiment 1, images (1936x1460 pixels, pixel size 0.23 µm) were obtained using APOTOME ZEISS Axioimager Z1 fluorescence microscope. Z-stacks (1 µm-spaced) were taken from brain sections of 4 mice/group/sex with a ZEISS Axiocam CCD monochrome camera and ZEN software (ZEISS Microscopy) at x20 magnification. For the experiment 2, images (2048x2048 pixels, pixel size 0.33 µm) were obtained using a Leica Thunder Tissue 3D widefield fluorescence microscope (PRIMACEN platform, Rouen, France). Z-stacks (0.5 µm-spaced) were taken from 6 mice/group/sex using a DFC9000 GT VSC-12292 monochrome camera with Las X Navigator software (Leica Microsystems) at x20 magnification. Identical illumination and exposure settings were applied for all images recorded. From the maximal intensity projection of these z-stacks, the GFAP- and IBA1 positive cells were counted manually and bilaterally by drawing region of interest (ROI) corresponding to the ARC of the hypothalamus using ImageJ software (National Institutes of Health, Bethesda, Maryland, US). The number of GFAP+ and IBA1+ cells was calculated in each ROI (1 per hemisphere) /mm² from 2-4 images per animal.

#### 2.4.2 Confocal microscopy and 3D IMARIS analysis

For the 3D reconstruction of microglia and astrocytes, images (1024x1024 pixels, pixel size 0.23 µm) were obtained using a TCS SP8 DM6000B-CFS confocal microscope with a Leica DFC 365 Fx camera (x63 oil-immersion lens, zoom 0.75). Z-stacks (0.3 µm steps) were taken in the ARC (1 image/hemisphere). All images were taken with the same confocal settings (pinhole, laser intensity, sequential mode). Raw files were then converted and analysed using IMARIS software (version 10.0.1, Oxford Instruments). First, the software was used to manually isolate cells using the cut function and to reconstruct the microglial surface using appropriate custom settings and to obtain volume object. Cells were included in the analysis when their soma was located within the central part of the z-stack and not cut by either the x or y plane (3-4 cells/hemisphere/animal). All morphological parameters were obtained after manually traced the IBA1 and GFAP staining using the Filament tracer mode. The volume as well as the number of total branch points and terminal points was measured for each cell. Branch Depth was defined as the number of branch points, or bifurcations, in the most complex path from the beginning point to a terminal point. Full Branch Level represented the highest value of Branching Level for the entire modeled cell. Filament Length corresponded to the sum of the lengths of all filaments. In addition, Sholl analysis was performed from the filament reconstruction mode.

### 2.5 RNA extraction and real-time quantitative polymerase chain reaction (RT-qPCR)

The hypothalamus was homogenized in 500 µL of liquid TRIzol Reagent (Invitrogen). Then, the homogenized samples were incubated for 5 min at RT and 100 µL of liquid chloroform (Merck Millipore, Fontenay Sous Bois, France) were added, followed by centrifugation at 12000*g* (15 min, 4°C). Following centrifugation, the aqueous phases containing the RNAs were transferred in new tubes and 250 µL of isopropanol was added to each tube. The samples were incubated for 10 min at RT, then centrifuged at 12000*g* at 4°C. The precipitated RNA formed a white pellet on the bottom of the tubes. The supernatant was removed, and the pellets were washed 3 times with 75% ethanol, followed at each step by centrifugation (8000*g*; 3 min; 4°C). The RNA pellets were air-dried before being resuspended in a volume of RNase-DNase free water (defined according to the size of the pellet obtained) for 1 hour on ice.

Tubes contained RNA were incubated at 65°C for 5 min. The total quantity and purity of the extracted RNA were determined by measuring the absorption at 260 and 280 nm using a Nanodrop 2000 spectrophotometer (Thermo Fisher Scientific).

For reverse transcription, each sample was diluted to obtain 1 μg of RNA in 8 μL of RNase-DNase free water. After adding 1 μL of 10X DNase Buffer and 1 μL of DNase (1 U/μL), the diluted samples were placed in a thermocycler (Eppendorf Mastercycler, Montesson, France) for 30 min at 37°C. The samples were incubated for 10 min at 65°C after the addition of 1 μL of DNase stop buffer to each tube. Then, 9 μL of the following mixture was added: 0.425 μL of Ribolock (40 U/μL, Thermo Scientific), 4 μL of 5X buffer, 1 μL of dNTP (10 mM), 0.5 μL of random hexamers (0.2 μg/μL), 2 μL of DTT (0.1 M), 1 μL of RT-MMLV enzyme (200 U/μL, Invitrogen), and 0.075 μL of H2O (Gibco) per tube. Finally, the samples were incubated at 42°C (50 min) followed by 5 min at 95°C.

Real-time PCR was performed using the Thermocycler Bio-Rad C1000 Touch Real-Time Thermal Cycler CFX96 (Bio-Rad). On a 96-well plate, 5 μL of SYBR Green mix (Bio-Rad) and 0.9 μL of RNase-DNase free water (Promega, Charbonière-les-Bains, France) were added to each well in addition to 0.05 μL of specific sense and antisense primers (100 mM, **Supplementary Table S2**). Then, 4 μL of diluted sample, range or RNase-DNase free water (negative control) were added into this mixture. Each cycle consists of a denaturation phase at 95°C, a hybridization phase whose temperature depends on each pair of primers (Tm) and a polymerization phase at 72°C. A mix of cDNA products was also prepared and then diluted to obtain a standard curve. The relative mRNA levels were assessed using *β-actin* and *Gapdh* as references and normalized by calculating the ratio of the starting quantity (SQ) of the gene of interest/mean (SQ household genes) provided by the Bio-Rad - CFX Manager 3.1 software (Bio-Rad Laboratories).

### 2.6 Statistics

All the statistical analyses were performed with GraphPad Prism version 8.0.1 (GraphPad Software La Jolla, CA). The ROUT method was used to identify outliers with a Q coefficient equal to 1%. Data of experiment 1 (w14) were compared with unpaired *t* tests or Mann-Whitney tests after assessing normality with the Shapiro-Wilk test. For data of experiment 2 (d3/d7/d14/d28), the effects of HFD and of the different time exposures were assessed using 2-way ANOVA (HFD x time) followed by Sidak’s multiple comparisons test. For the microscopy data, values were compared with nested *t* tests considering each subcolumn is for a replicate (animal) and each row in that subcolumn for a technical replicate (image or individual cell). For body composition and morphometric parameters, additional graphs display the normalization (HFD/SD) and values between male and female mice were compared with unpaired *t* tests. All graphs are presented as mean ± standard error of the mean (SEM) with the following significance levels: *p < 0.05, **p < 0.01, ***p < 0.001, ****p < 0.0001.

## 3. Results

### 3.1 HFD consumption led to similar body weight gains in male and female mice after 14 weeks

Male and female mice fed a high-fat diet (M- and F-HFD, respectively) gained more weight and exhibited more fat mass (in g and % of body weight) than control mice fed with a standard diet (SD) (*p*<0.0001, **Fig. 1A, B, C**) after 14 weeks (w14). More precisely, M-HFD and F-HFD gained in average 18.6 g and 13.1 g *vs* 8.1 g and 5.5 g for respective controls (M-SD and F-SD) and presented in average 11.9 g (29.3%) and 9.7 g (34%) *vs* 2.1 g (7.6%) and 2.2 g (11.5%) of fat mass (in g and % of body weight), respectively. There was no significant difference between the SD and HFD groups regarding lean mass (in g), although a strong trend to an increase was observed for the F-HFD group (*p*=0.0557, **Fig. 1D**). Nevertheless, the lean mass level (in % of body weight) was significantly decreased in both the M- and F-HFD groups (*p*<0.0001, **Fig. 1E**). Despite the ratio of body weight gain/initial weight of M-HFD was significantly higher from the w5 (p<0.01 *vs* M-SD, **Fig. 1F**) compared to only from the w7 for F-HFD (p<0.05 *vs* F- SD, **Fig. 1G**), the ratio was ultimately similar between the two sexes at w14 **(Fig. 1H)**.

**Fig. 1:**
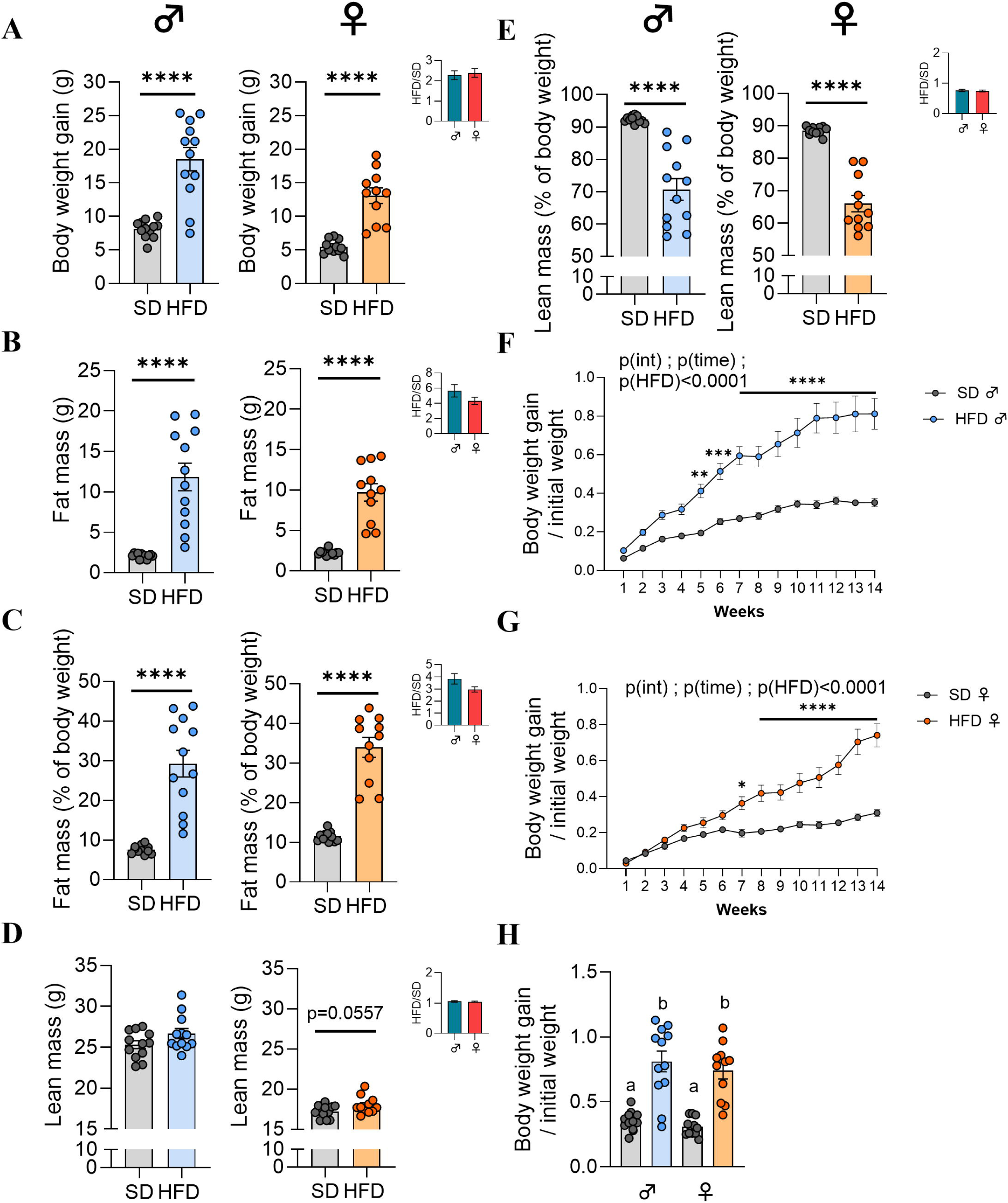
Monitoring of body weight and body composition after 14 weeks of high fat diet. (A) Body weight gain in grams (g), (B-E) fat mass and lean mass (in g and % of body weight) measured by EchoMRI in C57Bl/6J male and female mice fed either a standard diet control (SD) or high fat diet (HFD) for 14 weeks (experiment 1, n=12/group). Data were compared with unpaired *t* tests (\*\*\*\**p*<0.0001 *vs* SD). (F-H) Ratio of the body weight gain/initial weight. (F, G) Data were compared with 2-way ANOVA (HFD x time) followed by Sidak’s multiple comparisons test (*\*p*<0.05). (H) Data were compared with unpaired *t* tests (values without a common letter differ significantly). Additional graphs show the comparison between male and female mice for each parameter by calculating the fold changes for each HFD mouse compared to the mean of their respective control group (HFD/SD). Data were compared with unpaired *t* tests (\**p*<0.05) and are presented as mean ± standard error of the mean (SEM).

### 3.2 HFD female mice exhibited a more pronounced satiety profile than male mice after 14 weeks

The consumption of a hyperlipidic diet led to weight gain and greater fat storage in HFD-fed mice. Thus, we wanted to assess the effects of HFD feeding on signalling pathways involved in food intake and energy balance regulation. For that purpose, the levels of mRNA transcripts encoding orexigenic and anorexigenic neuropeptides were measured in the hypothalamus. The gene expression levels of neuropeptide Y (NPY) and agouti-related protein (AgRP) were downregulated both in M-HFD (*p*=0.0516 and *p*<0.01, respectively) and in F-HFD (*p*<0.01, **Fig. 2A, C**). Interestingly, the mRNA expression of the proopiomelanocortin (POMC) gene was only upregulated in F-HFD (*p*<0.05, **Fig. 2B**). Finally, the mRNA levels of the melanocortin-4 receptor (MC4R) in HFD groups were not significantly different from those in the control groups in either sex after w14 **(Fig. 2D)**.

**Fig. 2:**
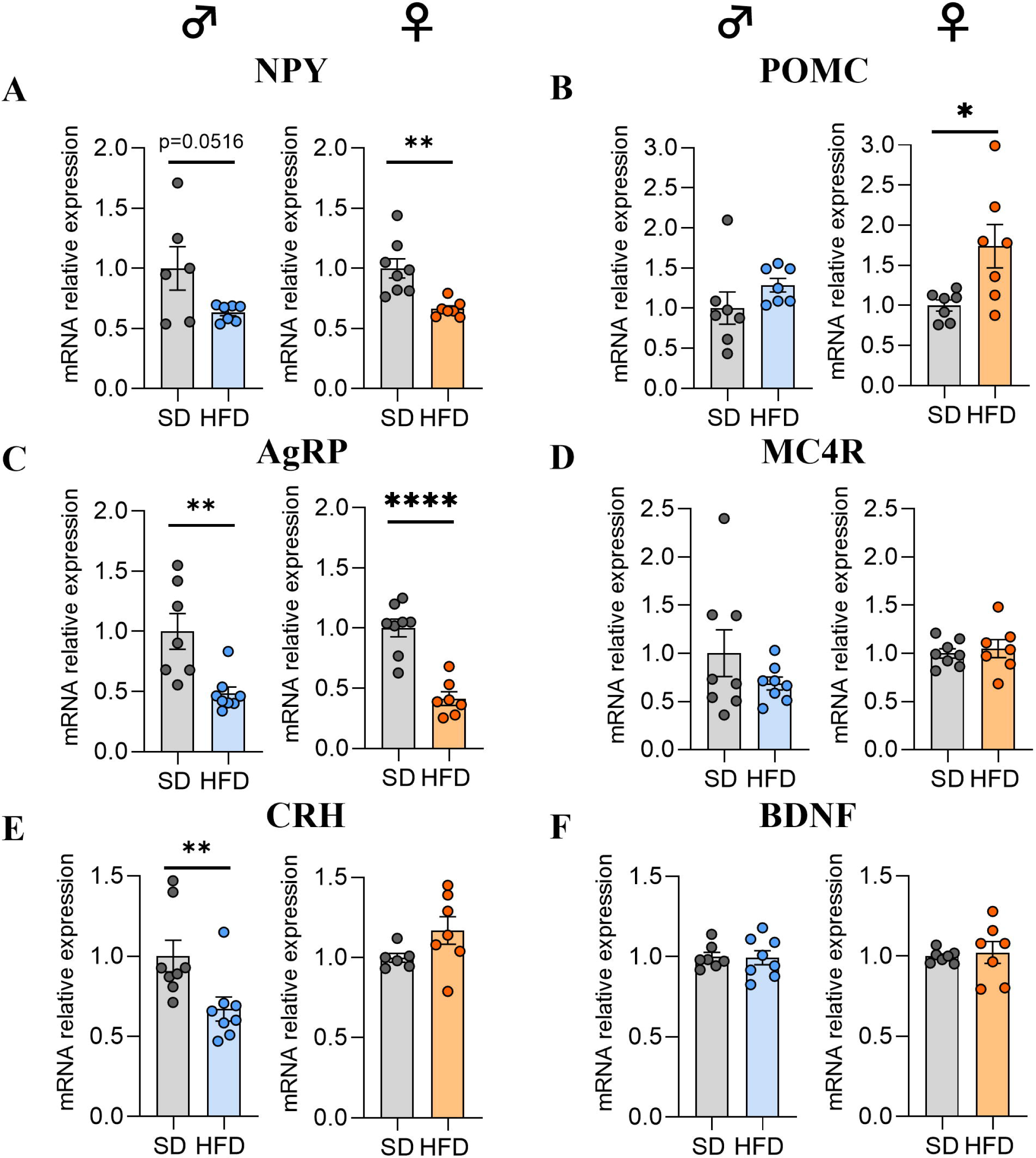
Effects of HFD for 14 weeks on hypothalamic mRNA expression of neuropeptides regulating food intake. (A-D) Relative quantification of mRNA transcript levels encoding orexigenic neuropeptides neuropeptide Y (NPY), agouti-related peptide (AgRP) and anorexigenic neuropeptides pro-opiomelanocortin (POMC) and melanocortin-4 receptor (MC4R) in hypothalamus of C57Bl/6J male and female mice fed for 14 weeks either SD or HFD (experiment 1, n=8/group). (E-F) Relative quantification of mRNAs transcript levels encoding the corticotropin-releasing hormone (CRH) and brain-derived neurotrophic factor (BDNF) in hypothalamus of C57Bl/6J male and female mice fed for 14 weeks either SD or HFD (experiment 1, n=8/group). All mRNA levels were quantified relative to *Gapdh* and *β-actin* housekeeping gene expressions and compared with unpaired *t* tests or Mann-Whitney tests (\**p*<0.05, \*\**p*<0.01, \*\*\*\**p*<0.0001 *vs* SD). All graphs show the fold changes of mRNA levels for each HFD mouse compared to the mean of their respective control group. Data are presented as mean ± standard error of the mean (SEM).

### 3.3 HFD male mice showed lower CRH gene expression levels after 14 weeks of HFD feeding

Corticotropin-releasing hormone (CRH) is known to coordinate the behavioural stress response (Hu et al., 2020) and is also involved in the control of energy homeostasis (Schwartz et al., 2000). Thus, we were interested in the effects of HFD consumption on the gene expression regulation of this peptide. Interestingly, mRNA levels of CRH were only decreased in M-HFD after w14 (*p*<0.01, **Fig. 2E**). Because its expression could be regulated by nutritional state and by MC4R signalling (Xu et al., 2003), we also analysed the mRNA transcript encoding brain-derived neurotrophic factor (BDNF). No significant difference was observed between the SD and HFD groups for either sex (**Fig. 2F**).

### 3.4 HFD consumption did not induce a major hypothalamic inflammatory response in male or female mice after 14 weeks

After studying the expression profile of major hypothalamic neuropeptides, we next investigated the impact of a HFD on hypothalamic inflammatory markers. First, we analysed the mRNA expression levels encoding proinflammatory cytokines. Overall, no significant changes were found in the interleukin-1β (IL-1β), interleukin-6 (IL-6) or α-tumor necrosis factor (TNF-α) levels in both sexes (**Fig. 3A, B, C**).

**Fig. 3:**
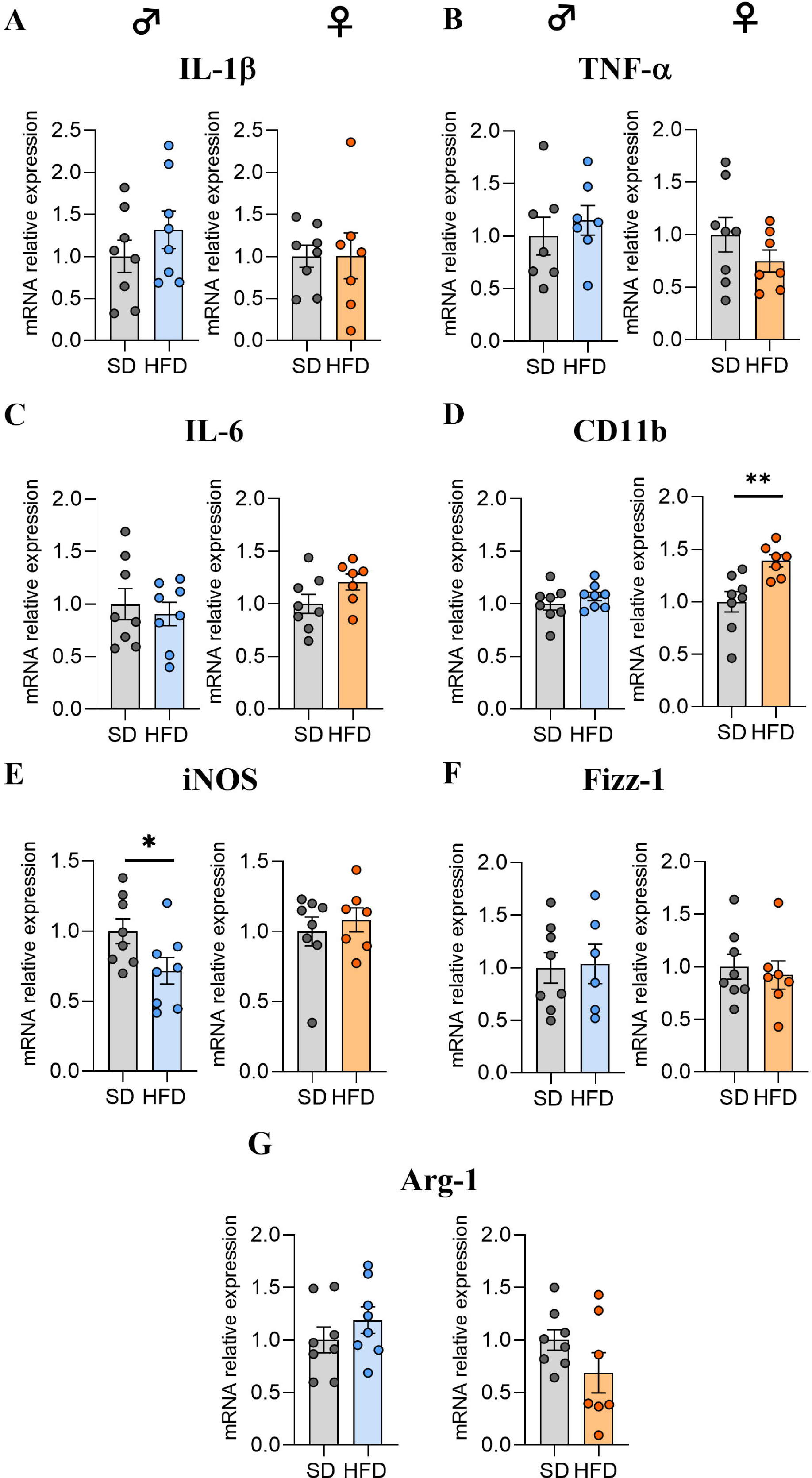
Effects of HFD feeding for 14 weeks on inflammatory marker expression levels in hypothalamus. (A-C) Relative quantification of mRNA transcript levels encoding proinflammatory cytokines with interleukin-1β (IL-1β), α-tumor necrosis factor (TNF-α), and interleukin-6 (IL-6) in hypothalamus of C57Bl/6J male and female mice fed for 14 weeks either SD or HFD (experiment 1, n=8/group). (D-G) Relative quantification of mRNA transcript levels encoding M1 polarization markers of macrophages (CD11b, iNOS) and M2 polarization markers (Fizz-1, Arginase-1) in hypothalamus of male and female C57Bl/6J mice fed for 14 weeks either SD or HFD (experiment 1, n=8/group). All mRNA levels were quantified relative to *Gapdh* and *β-actin* housekeeping gene expressions and compared with unpaired *t*-tests or Mann-Whitney tests (\**p*<0.05 *vs* SD). All graphs show the fold changes of mRNA levels for each HFD mouse compared to the mean of their respective control group. Data are presented as mean ± standard error of the mean (SEM).

To continue, we were interested in gene expression profiling of macrophages by targeting pro- and anti-inflammatory phenotypes with the expression of M1 (CD11b, iNOS) and M2 (Fizz-1, Arginase-1) polarization markers. Interestingly, the mRNA level of the gene encoding CD11b was only increased in F-HFD (*p*<0.01, **Fig. 3D**), while that of iNOS was only decreased in M-HFD (*p*<0.05, **Fig. 3E**). However, no significant changes in Fizz-1 or Arginase-1 levels were found in male or female mice (**Fig. 3F, G**).

### 3.5 Long-term HFD exposure induced the downregulation of IBA1 in female mice

According to the literature, a series of cellular and molecular events involving microglial cells and astrocytes, collectively termed “reactive gliosis”, are observed in rodents fed with a HFD (Lee et al. 2020). Therefore, we investigated the glial response by first performing RT‒qPCR of mRNA transcripts of genes classically associated with microgliosis and astrogliosis, such as the ionized calcium-binding adaptor molecule 1 (IBA1) protein, the purinergic receptor P2Y12 (P2RY12) and the glial fibrillary acidic protein (GFAP). HFD exposure for 14 weeks only led to a decrease in the gene expression of the microglial marker IBA1 in the hypothalamus as well as the IBA1/P2RY12 ratio in F-HFD (*p*<0.05, **Fig. 4A, B**).

**Fig. 4:**
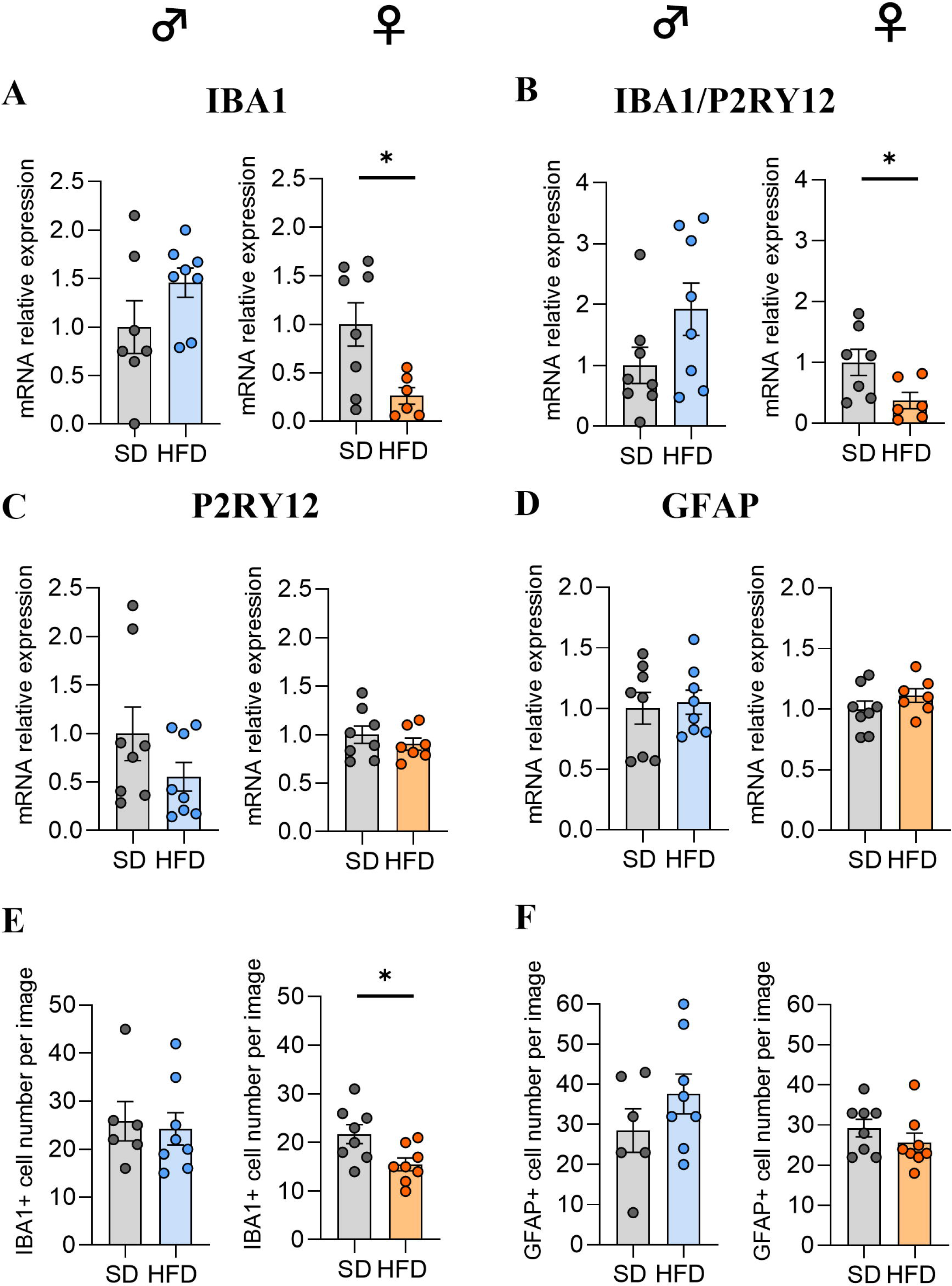
Effects of HFD feeding for 14 weeks on glial cell marker expression levels in hypothalamus. (A-D) Relative quantification of mRNA transcript levels encoding microglial markers with ionized calcium-binding adapter molecule 1 (IBA1), purinergic receptor P2Y12 (P2RY12) and astrocytic marker glial fibrillary acidic protein (GFAP) in hypothalamus of C57Bl/6J male and female mice fed for 14 weeks either SD or HFD (experiment 1, n=8/group). All mRNA levels were quantified relative to *Gapdh* and β*-actin* housekeeping gene expressions and compared with unpaired *t*-tests or Mann-Whitney tests (\**p*<0.05 *vs* SD). All graphs show the fold changes of mRNA levels for each HFD mouse compared to the mean of their respective control group. (E-F) Quantification of detection by immunofluorescence of IBA1 and GFAP proteins within the ARC from C57Bl/6J male and female mice fed for 14 weeks either SD or HFD (experiment 1, N=2 images/animal with n=4 mice/group). Immunopositive cells for IBA1 and GFAP were manually and bilaterally counted using Image J software in coronal sections of the ARC (20 µm, -1.22 to 2.54 mm relative to Bregma). Data are presented as mean ± standard error of the mean (SEM). Values were compared with nested *t* tests (\**p*<0.05 *vs* SD).

This quantification of multiple markers *via* RT‒qPCR was complemented by histological analysis with immunofluorescence staining of astrocytes and microglial cells in the ARC. Interestingly, the genetic analysis described above have been corroborated with the preliminary counting which displayed a decrease of the number of IBA1 positive cells in the F-HFD group (p<0.05, **Fig. 4E, Supplementary Fig. S1**). Similar to the results of the GFAP gene expression analysis, the numbers of GFAP+ cells in the ARC were equivalent between the SD-fed and HFD-fed mice (**Fig. 4F, Supplementary Fig. S1**).

These preliminary data did not seem to support the presence of hypothalamic inflammation but showed a sex-divergent response after long-term HFD exposure. Several studies have reported that hypothalamic inflammation occurs very early in response to a HFD before the onset of obesity (Cansell et al., 2021; Thaler et al., 2012). Therefore, we studied the hypothalamic response in a second cohort of male and female mice on a shorter timescale by monitoring the effects of 3, 7, 14 and 28 days of exposure to a HFD and performed the same analyses within the hypothalamus.

### 3.6 HFD male mice gained more weight, body fat and more rapidly than HFD female mice during a short-term exposure

Overall, HFD consumption affected body weight gain (*p(HFD)*<0.001, **Fig. 5**), fat mass (in g and %) (*p(HFD)*<0.01, **Fig. 6A-D**) and lean mass percentage (*p(HFD)*<0.001, **Fig. 6G, H**) in male and female mice. In addition, there was also a time exposure effect on body weight gain (*p(time)*<0.0001, **Fig. 5**) and on lean mass (in g and %) for both sexes (*p(time)*<0.01, **Fig. 6E-H**) and on fat mass (in g and %) but only for male mice (*p(time)*<0.0001, **Fig. 6A, C**).

**Fig. 5:**
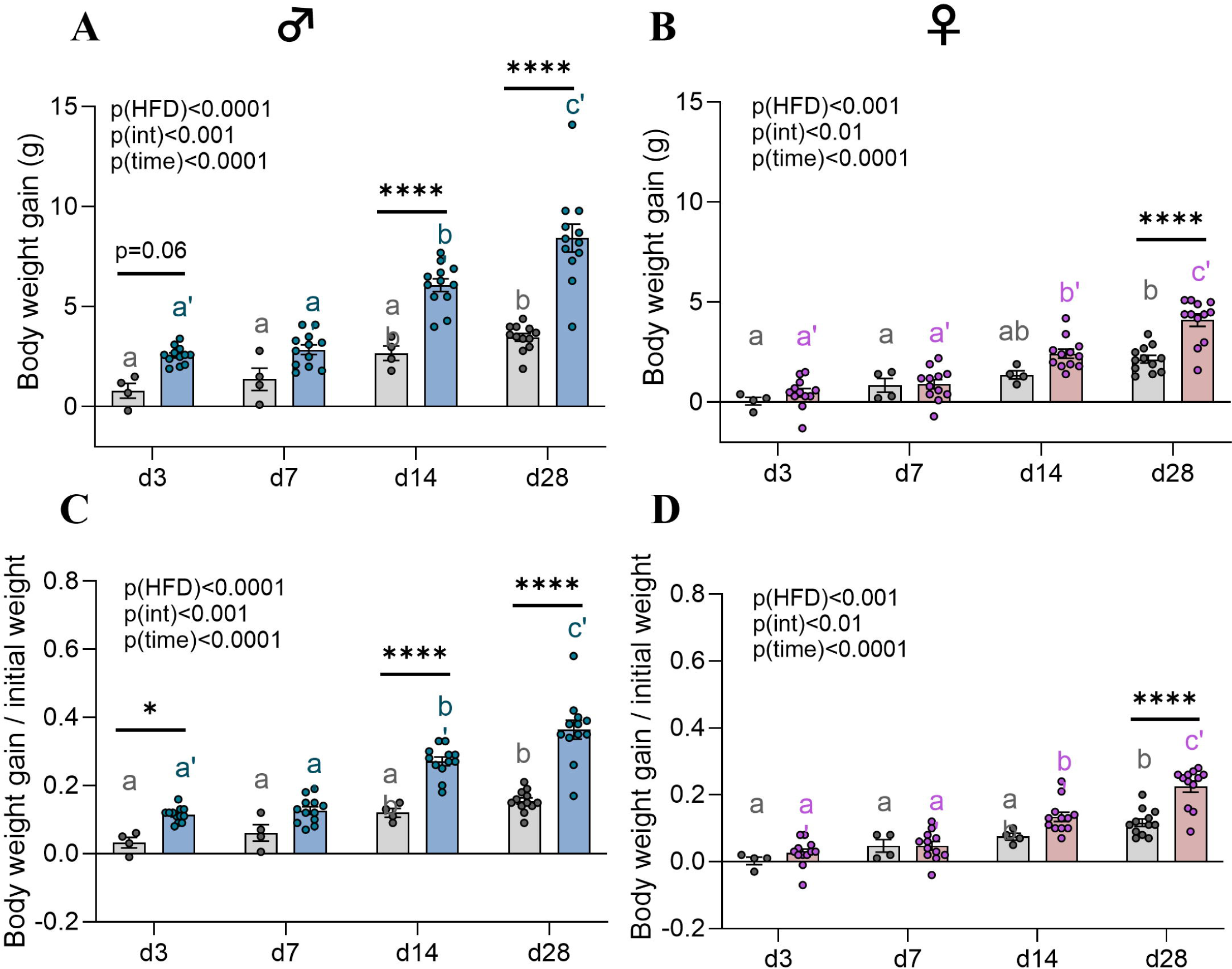
Body weight gain after 3, 7, 14 and 28 days of HFD feeding. (A, B) Body weight gain in grams (g) and (C, D) ratio of the body weight gain/initial weight in C57Bl/6J male and female mice fed either SD or HFD feeding for up 28 days (experiment 2, n=4 SD for d3, d7, d14; n=12 SD for d28 and n=12 HFD/time). Values were compared with 2-way ANOVA (HFD x time) followed by Sidak’s multiple comparisons test: HFD *vs* SD groups/time (\**p*<0.05, \*\**p*<0.01, \*\*\**p*<0.001, \*\*\*\**p*<0.0001), between SD groups (a,b,c letters) and between HFD groups (a’,b’,c’ letters). Values without a common letter differ significantly. Data are presented as mean ± standard error of the mean (SEM).

**Fig. 6:**
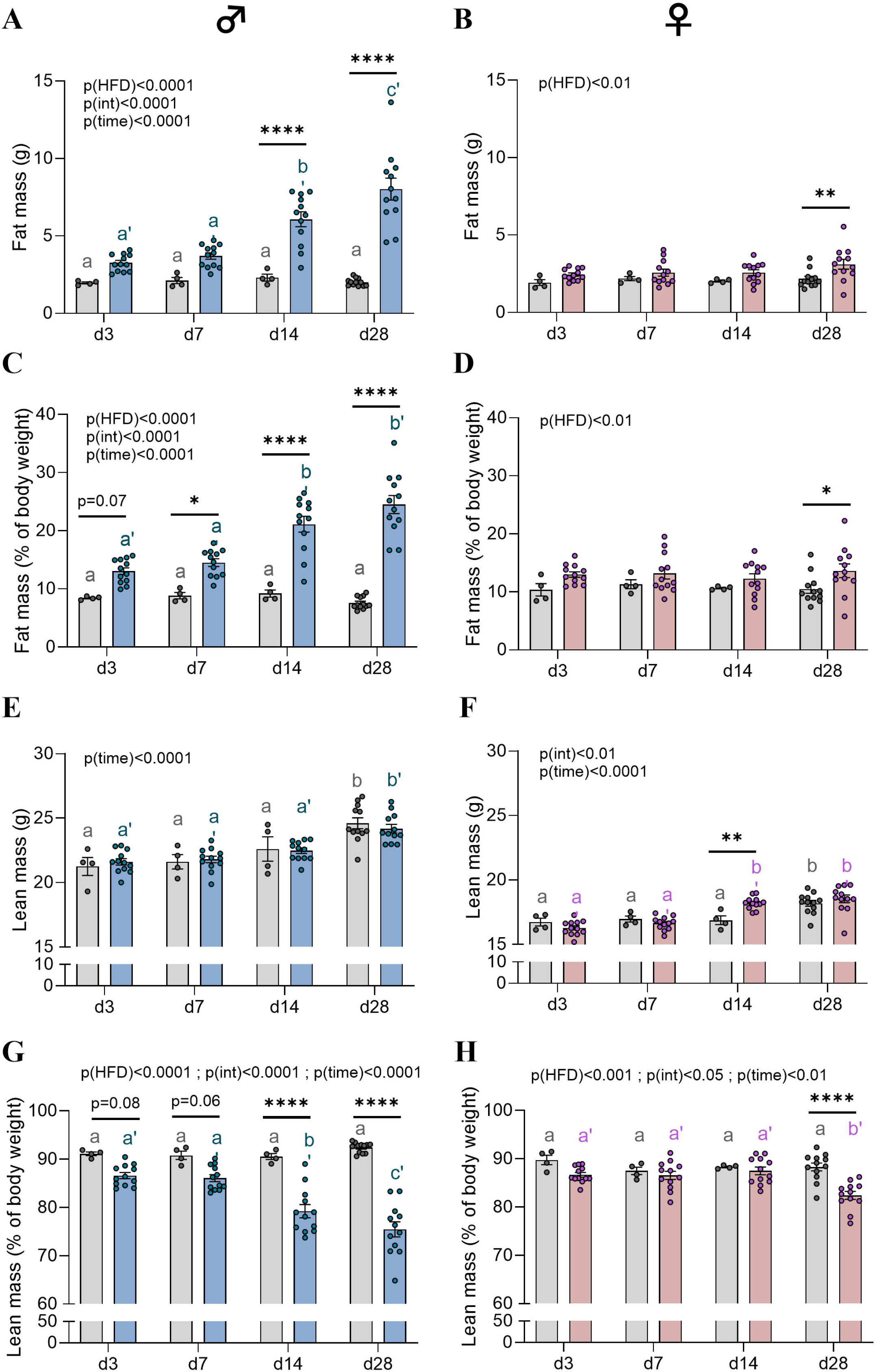
Body composition changes after 3, 7, 14 and 28 days of HFD feeding. (A-D) Fat mass and (E-H) lean mass (in g and % of body weight) measured by EchoMRI in C57Bl/6J male and female mice fed either SD or HFD feeding for up 28 days (experiment 2, n=4 SD for d3, d7, d14; n=12 SD for d28 and n=12 HFD/time). Values were compared with 2-way ANOVA (HFD x time) followed by Sidak’s multiple comparisons test: HFD *vs* SD groups/time (\**p*<0.05, \*\**p*<0.01, \*\*\**p*<0.001, \*\*\*\**p*<0.0001), between SD groups (a, b, c letters) and between HFD groups (a’, b’, c’ letters). Values without a common letter differ significantly. Data are presented as mean ± standard error of the mean (SEM).

More precisely, the body weight gain (in g) in male mice was not significant after 3 days (d3) of HFD feeding despite a strong trend (*p*=0.06 *vs* M-SD, **Fig. 5A**), but it was starting from d14 to d28 (*p*<0.0001 *vs* M-SD, **Fig. 5A**) and at d3 in regard to their initial weight (*p*<0.05 *vs* M-SD, **Fig. 5C**). In contrast, female mice showed significant weight gain only from d28 of HFD (*p*<0.0001 *vs* F-SD, **Fig. 5B, D**). On average, M-HFD gained 2.6 g, 2.9 g, 6 g and 8.4 g compared to 0.05 g, 0.9 g, 2.4 g, and 4.1 g for F-HFD after d3, d7, d14 and d28, respectively **(Fig. 5)**.

M-HFD-fed animals displayed an increase in fat mass at earlier time points than F-HFD-fed mice. Indeed, the fat mass (in g) was significantly greater in M-HFD mice starting from d14 to d28 (*p*<0.0001 *vs* M-SD, **Fig. 6A**) than in F-HFD mice from d28 (p<0.01 *vs* F-SD, **Fig. 6B**). When expressed as % of body weight, the fat mass increase was trending at d3 (*p*=0.07 *vs* M-SD) and was significant starting from d7 (*p*<0.05 *vs* M-SD) to d28 (*p*<0.0001 *vs* M-SD, **Fig. 6C**) in the M-HFD groups. The increase in the F-HFD group was only significant at d28 (*p*<0.05 *vs* F-SD, **Fig. 6D)**. Specifically, M-HFD exhibited greater fat mass than did F-HFD with in average fat masses of 3.2 g (13.03%), 3.7 g (14.5%), 6 g (21.1%) and 8 g (24.5%) compared to 2.4 g (12%), 2.6 g (13.2%), 2.6 g (12.3%), and 3.1 g (13.6%) for F-HFD at d3, d7, d14 and d28, respectively (**Fig. 6A-D**). On the one hand, the absolute lean mass (g) was not significantly different between the M-SD and M-HFD groups (**Fig. 6E**), but it was between M- and F-HFD at d3 (p<0.05) and d14 (p<0.0001, **Supplementary Fig. S2C**). On the other hand, the proportion of lean mass (in %) tended to decrease for M-HFD at d3 (*p*=0.08 *vs* M-SD) and d7 (*p*=0.06 *vs* M-SD), and it was significantly decreased at d14 and d28 (*p*<0.0001 *vs* M-SD, **Fig. 6G**) and only at d28 for F-HFD (*p*<0.0001 *vs* F-SD, **Fig. 6H**). Interestingly, F-HFD exhibited a significant rise of lean mass (in g) compared to F-SD at d14 (*p*<0.01, **Fig. 6F**). This increase was not observed in M-HFD. In addition, female HFD/SD ratio was higher at d14 and from d7 (in g and %, respectively) compared to males (p<0.05, Supplementary **Fig. S2C, D**).

### 3.7 Short-term HFD exposure led to specific modulation of neuropeptide gene expression involved in energy balance regulation in a time-dependent manner between male and female mice

Consistently with the first cohort (experiment 1), the expression of mRNAs encoding NPY and AgRP were also decreased in male and female mice after short-term HFD consumption (*p(HFD)*<0.0001, *p(time)*<0.01; **Fig. 7A, C)**. More precisely, mRNA levels encoding NPY tended to decrease at d3 (*p*<0.01) and d7 (p=0.0753 *vs* M-SD, p=0.0574 *vs* F-SD) for M- and F-HFD, respectively, but decreased significantly at d28 (*p*<0.001, **Fig. 7A**) only for M-HFD. Moreover, AgRP levels were also significantly decreased at d3 (*p*<0.001, **Fig. 7C**) for both sexes and at d7 and d28 but only for F-HFD *(p*<0.05 *vs* F-SD, **Fig. 7C**). The upregulation of POMC gene expression found in F-HFD mice at w14 (**Fig. 2B**) was transient at d14 (*p*<0.05 *vs* F*-*SD, **Fig. 7B**) but also in M-HFD, especially at d3 (*p*<0.01 *vs* M*-*SD, **Fig. 7B**). While HFD consumption did not affect the mRNA levels of the gene encoding CRH in both sexes in contrast to what was observed in M-HFD after w14 (**Fig. 2E**), interaction and time effects (*p<*0.05) were found for MC4R and BDNF factors, respectively, in female mice (**Fig. 7D, F**).

**Fig. 7:**
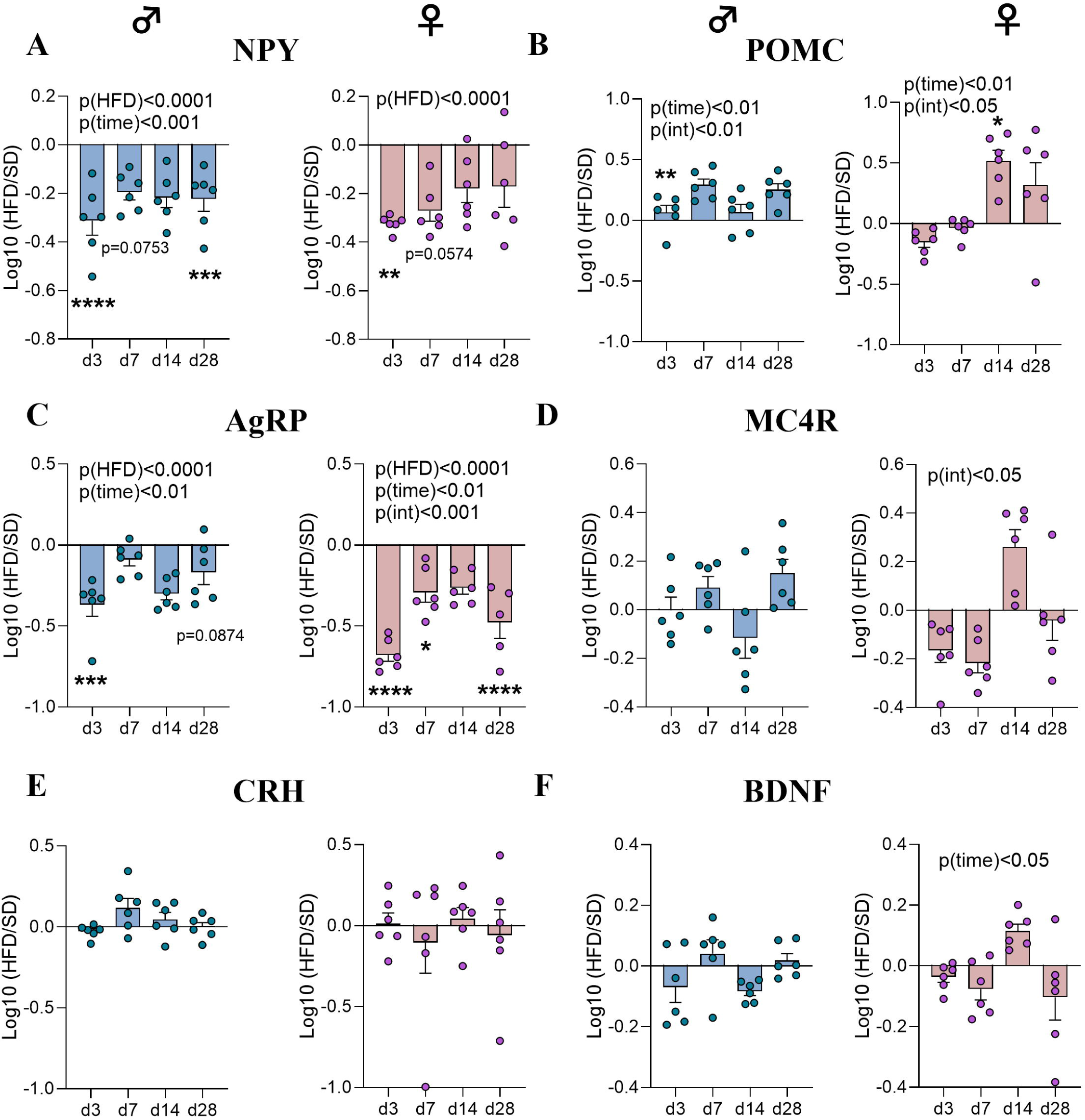
Time course response to HFD of hypothalamic neuropeptide gene expressions involved in food intake regulation. (A-D) Relative quantification of mRNA transcript levels encoding orexigenic neuropeptides with neuropeptide Y (NPY), agouti-related peptide (AgRP) and anorexigenic neuropeptides with pro-opiomelanocortin (POMC) and melanocortin-4 receptor (MC4R) in hypothalamus of male and female C57Bl/6J mice fed either SD or HFD for 3, 7, 14 and 28 days (experiment 2, n=6/group). (E-F) Relative quantification of mRNA transcript levels encoding the corticotropin-releasing hormone (CRH) and brain-derived neurotrophic factor (BDNF) in hypothalamus of C57Bl/6J male and female mice fed for 3, 7, 14 and 28 days (experiment 2, n=6/group). All mRNA species were quantified relative to *Gapdh* and *β-actin* housekeeping gene expression. Raw data were compared with 2-way ANOVA (HFD x time) followed by Sidak’s multiple comparisons test: HFD *vs* SD groups/time (\**p*<0.05, \*\**p*<0.01, \*\*\**p*<0.001, \*\*\*\**p*<0.0001). All graphs show the fold changes of mRNA levels for each HFD mouse compared to the mean of their respective control group (Log10 (HFD/SD)).

### 3.8 The HFD exposure was associated with earlier expression of inflammatory markers in female than male mice

As observed in the first experiment, mRNA levels of proinflammatory cytokines in HFD-fed mice were not significantly different from those in SD mice for either sex despite a trend toward an increase in IL-6 mRNA expression up to d28 in M-HFD (*p(HFD)*=0.0895, **Fig. 8C**). Nevertheless, the upregulation of CD11b observed at w14 in F-HFD (**Fig. 3D**) also occurred from d14 (*p(HFD)*=0.0587, *p(time)*=0.0588; **Fig. 8D**). In opposite, these female mice also exhibited downregulated iNOS gene expression (*p(time)=*0.0554, **Fig. 8E**) and Fizz-1 levels (*p(HFD*)=0.09, *p(int)* and *p(time)*<0.01, **Fig. 8F**), specifically at d3 (*p*<0.001 *vs* F-SD, **Fig. 8F**), as did M-HFD-fed mice but at d7 *(p(int)<0.05; p*<0.05 *vs* M-SD, **Fig. 8F**).

**Fig. 8:**
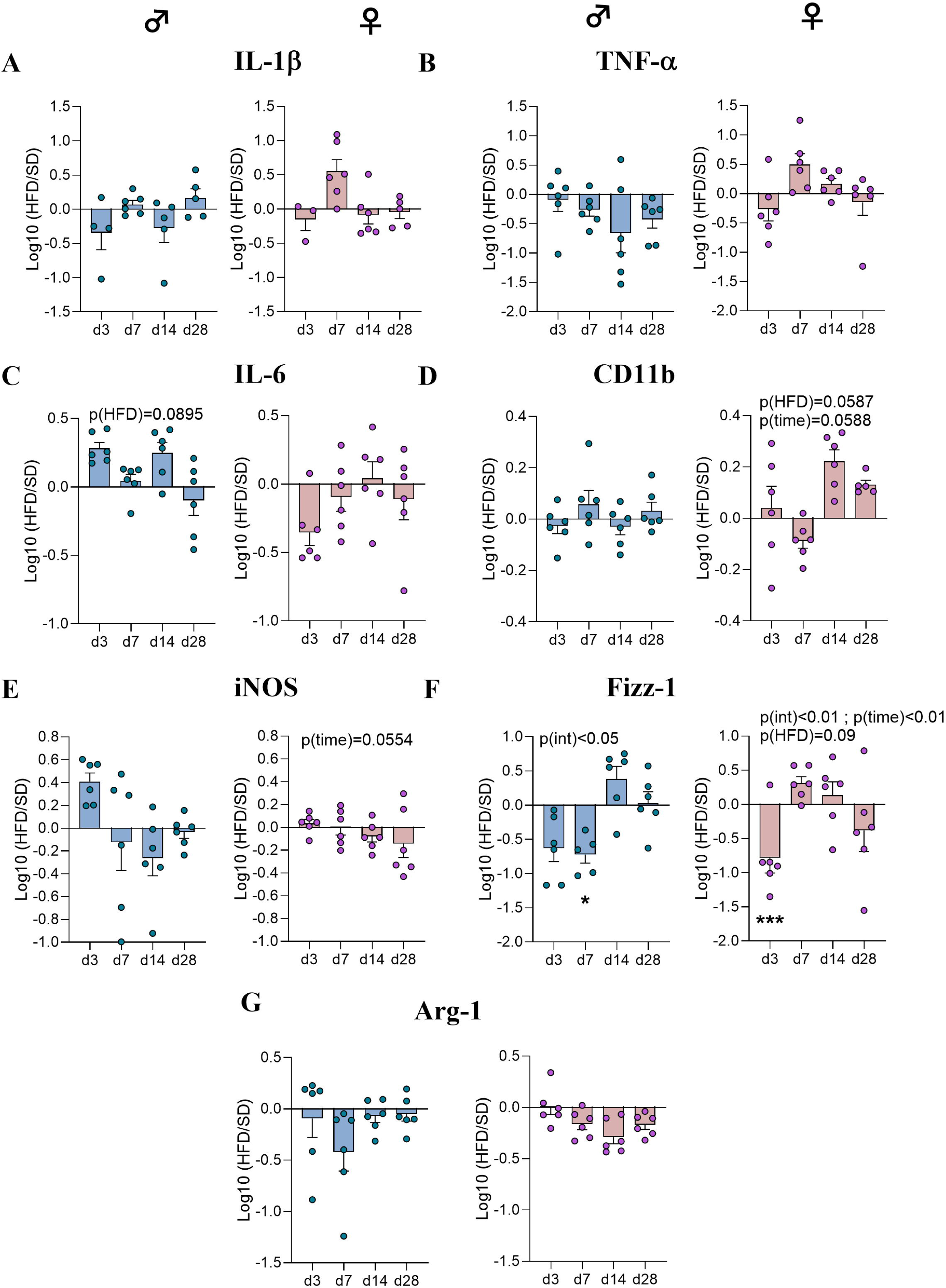
Time course response to HFD of hypothalamic inflammatory markers. (A-C) Relative quantification of mRNAs transcript levels encoding proinflammatory cytokines with interleukin-1β (IL-1β), α-tumor necrosis factor (TNF-α), interleukin-6 (IL-6) in hypothalamus of C57Bl/6J male and female mice fed either SD or HFD for 3, 7, 14 and 28 days (experiment 2, n=6/group). (D-G) Relative quantification of mRNAs transcript levels encoding M1 polarization markers of macrophages (CD11b, iNOS) and M2 polarization markers (Fizz-1, Arginase-1) in hypothalamus of C57Bl/6J male and female mice fed either SD or HFD for 3, 7, 14 and 28 days (Experiment 2, n=6/group). All mRNA species were quantified relative to *Gapdh* and *β-actin* housekeeping gene expression. Raw data were compared with 2-way ANOVA (HFD x time) followed by Sidak’s multiple comparisons test: HFD *vs* SD groups/time (*p<0.05, **p<0.01, ***p<0.001, ****p<0.0001 *vs* SD). All graphs show the fold changes of mRNA levels for each HFD mouse compared to the mean of their respective control group (Log10 (HFD/SD)).

### 3.9 Short-term HFD exposure did not induce relevant changes in genes related to glial markers in male and female mice

In contrast to previous studies (Thaler et al., 2012) and although a decrease in IBA1 expression occurred in F-HFD at w14 (**Fig. 4A**), we did not observe significant effects of HFD on IBA1 gene expression at earlier time points in male or female mice (data not shown). However, a time-dependent downregulation of GFAP gene expression was observed in both sexes (p(*time*)<0.01, **Fig. 9A**). Our immunofluorescence analyses revealed a decreasing trend in the number of IBA1+ cells in the ARC of M-HFD at d7 (p=0.0880 *vs* M-SD, **Fig. 9B**). Interestingly, we also reported trends among M-HFD mice, with a transient increase at d14 compared to those at d7 (p=0.0685) and d28 (p=0.07, **Fig. 9B**). In addition, we found a significant decrease in the number of GFAP+ cells in the F-HFD group at d3 (p<0.01, **Fig. 9C**).

**Fig. 9:**
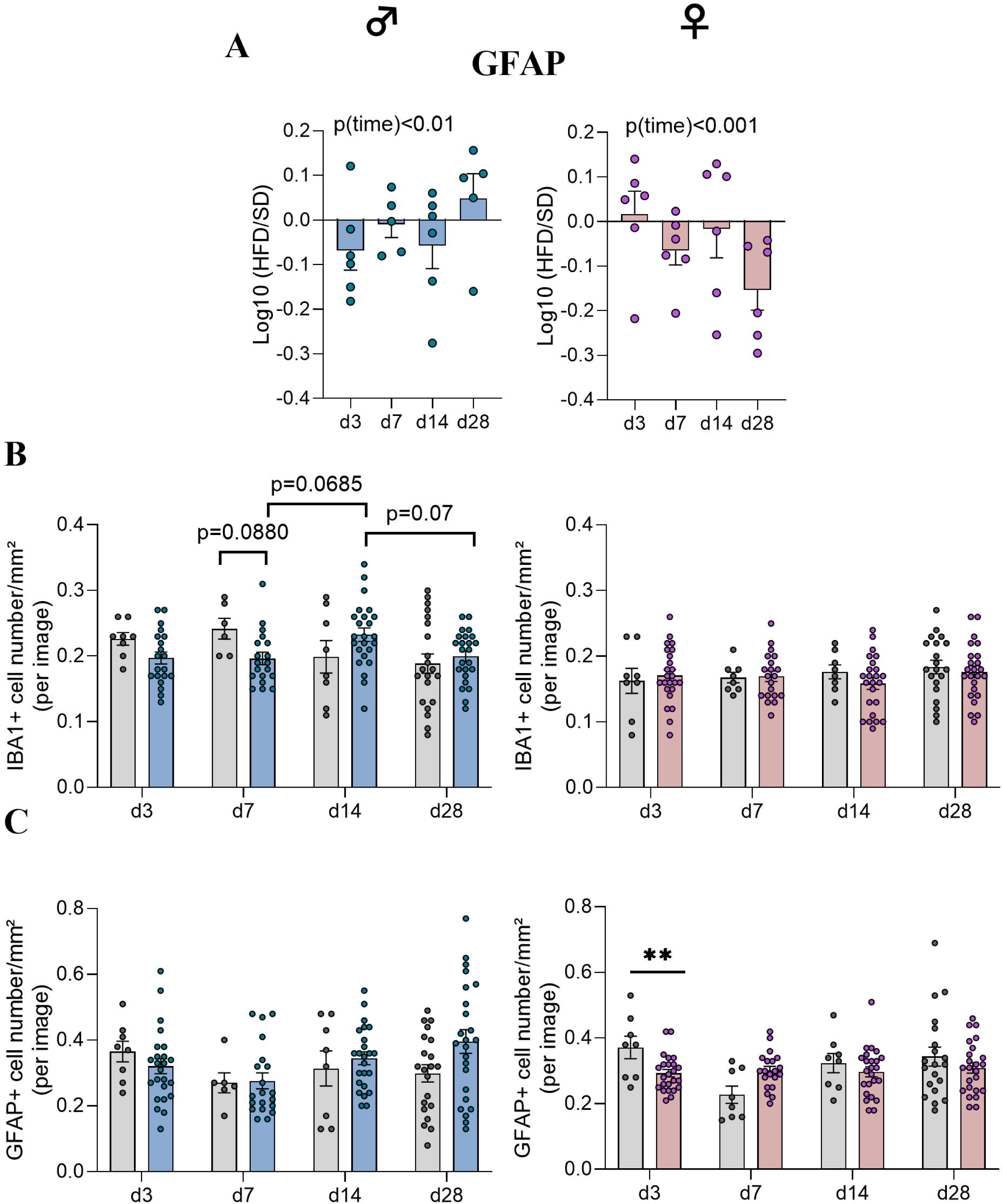
Time course response to HFD of glial marker levels in hypothalamus. (A) Relative quantification of mRNA transcript levels encoding astrocytic marker glial fibrillary acidic protein (GFAP) in hypothalamus of C57Bl/6J male and female mice fed either SD or HFD for 3, 7, 14 and 28 days (experiment 2, n=6/group). All mRNA species were quantified relative to *Gapdh* and *β-actin* housekeeping gene expression. Raw data were compared with 2-way ANOVA (HFD x time) followed by Sidak’s multiple comparisons test: HFD *vs* SD groups/time (\**p*<0.05, \*\**p*<0.01, \*\*\**p*<0.001, \*\*\*\**p*<0.0001 *vs* SD). Graphs show the fold changes of mRNA levels for each HFD mouse compared to the mean of their respective control group (Log10(HFD/SD)). (B, C) Quantification of detection by immunofluorescence of IBA1 and GFAP proteins within the ARC from C57Bl/6J male and female mice fed for 3,7,14 and 28 days either SD or HFD (N= 4 images/animal with n=6 mice/group). Immunopositive cells for IBA1 and GFAP were manually and bilaterally counted using Image J software in coronal sections of the ARC (20 µm, -1.22 to 2.54 mm relative to Bregma). Values were compared with nested *t* tests: HFD *vs* SD / time, between HFD groups and between SD groups (*\*p*<0.05). Data are presented as mean ± standard error of the mean (SEM).

### 3.10 Signs of structural remodelling of microglia and astrocytes differed between male and female mice after 28 days and 14 weeks of HFD feeding

To further assess the glial response, we implemented three-dimensional morphometric analyses of individual microglial cells and astrocytes in the ARC at d28 and w14 after HFD feeding. Using IMARIS software, we assessed various features, including cell volume, glial branch parameters (filament length, filament full branch depth/level, and filament number segment branch/terminal points) and Sholl intersections. Interestingly, we firstly reported time-dependent remodelling of microglia in SD mice. More specifically, M-SD mice at w14 displayed microglia with more segment branch points (p=0.0544) and terminal points (p<0.05) than M-SD mice at d28 (**Fig 10B**). Conversely, microglia within F-SD at w14 displayed fewer segment branch/terminal points and filament full branch depth than did those within the F-SD group at d28 (p<0.05, **Fig. 10B**). A comparison between male and female mice revealed a lower cell volume (p<0.01, **Fig. 10B**), filament No. segment branch /terminal pts (p<0.05, Fig. **10B**), and branch depth (p=0.07, **Fig. 10B**) in F-HFD compared to M-HFD at d28. Moreover, we observed a lower filament length sum in the M-HFD group compared to the F-HFD group at w14 (p<0.05) and to the M-HFD group at d28 (p<0.01, **Fig. 10B**). Second, we found significant morphometric changes in astrocytes, such as an increase in the total filament volume and the number of segment branch points in the M-HFD and F-HFD groups at d28, respectively (p<0.05 *vs* SD, **Fig. 11B**). In addition, F-HFD showed a strong trend toward astrocytes with a greater number of segment terminal points (p=0.0640 *vs* F-SD) and filament length sum at d28 (p=0.0528 *vs* F-SD, **Fig. 11B**). By comparing male and female animals, we found a significant decrease in the sum of the filament volume in the F-HFD group compared to that in the M-HFD group at d28 (p<0.05, **Fig. 11B**).

**Fig. 10:**
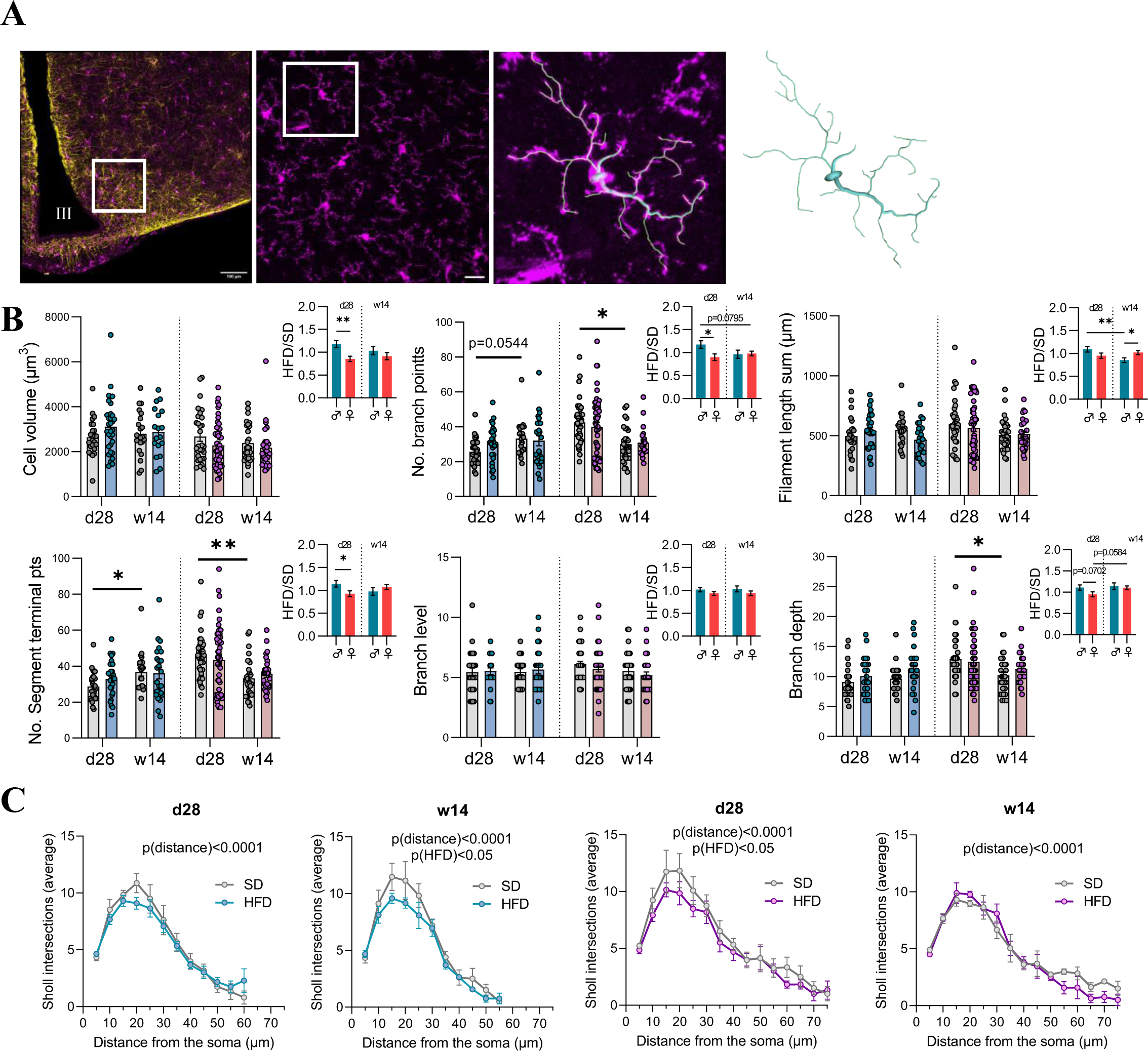
Morphometric analysis of microglia after 28 days and 14 weeks of HFD within the ARC. (A) First panel: representative image of a 3D mosaic (512x512 pixels, voxel size 0.459µm) from confocal microscope showing IBA1, GFAP and merge stained microglia (purple) and astrocytes (yellow) within the ARC. Scale bar: 100µm, III: third ventricle. Second panel: representative image showing IBA1 staining obtained after maximal projection Z-stacks (1024x1024 pixels, 234.55x234.55µm). Scale bar: 20µm. Third panel: focus of the white square drawn in the second panel which highlights an example of the filament tracing performed in isolated microglia. (B) Graphs show the values of each individual microglia for cell volume, Filament No. segment branch points/terminal points, filament full branch depth/level, and filament length sum for SD mice (grey bars) and HFD mice (blue and pink bars for male and female mice, respectively) at d28 (N=3-4 cells/hemi-ARC from n=6 mice/group) and w14 (N=3-4 cells/hemi-ARC from n=4 mice/group). Values were compared with nested *t* tests (HFD *vs* SD, between HFD groups and between SD groups*, *p*<0.05). Data are presented as mean±standard error of the mean (SEM). Additional graphs show the comparison between male and female mice for each parameter by calculating the fold changes for each HFD mouse compared to the mean of their respective control group (HFD/SD) at d28 and w14. Data were compared with unpaired *t* tests (*\*p*<0.05) and are presented as mean±standard error of the mean (SEM). (C) Graphs show the mean distribution of the number of Sholl intersections as a function of the distance from the microglial soma for SD mice (grey curves) and HFD mice (blue and pink curves for male and female mice, respectively) at d28 (N=3-4 cells/hemi-ARC from 6 mice/group) and at 14 weeks (N= 3-4 cells/hemi-ARC from 4 mice/group). Values were compared with 2-way ANOVA (HFD x distance) followed by Sidak’s multiple comparisons test (between HFD groups and their respective control group, *\*p*<0.05).

**Fig. 11:**
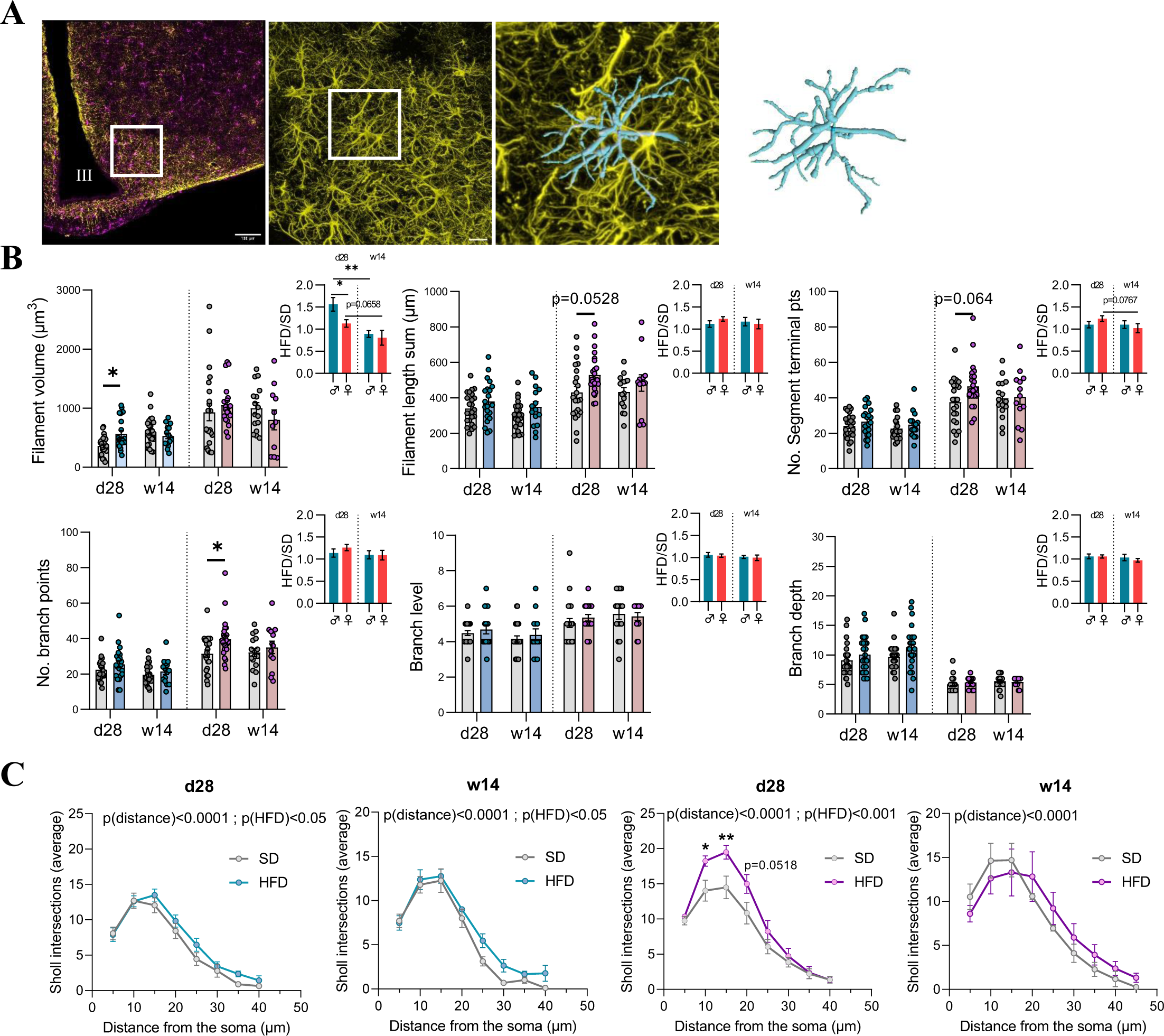
Morphometric analysis of astrocytes after 28 days and 14 weeks of HFD within the ARC. (A) First panel: representative image of a 3D mosaic (512x512 pixels, voxel size 0.459 µm) from confocal microscope showing IBA1, GFAP and merge stained microglia (purple) and astrocytes (yellow) within the ARC. Scale bar: 100 µm, III: third ventricle. Second panel: representative image showing GFAP staining obtained after maximal projection Z-stacks (1024x1024 pixels, 234.55x234.55 µm). Scale bar: 20 µm. Third panel: focus of the white square drawn in the second panel highlights an example of the surface reconstruction in isolated astrocytes. (B) Graphs show the values of each individual astrocytes for Filament volume sum, Filament No. segment branch points/terminal points, filament full branch depth/level, and filament length sum for SD mice (grey bars) and HFD mice (blue and pink bars for male and female mice, respectively) at d28 (N=3-4 cells/hemi-ARC from n=6 mice/group) and at w14 (N=3-4 cells/hemi-ARC from n=4 mice/group). Values were compared with nested *t*-tests (HFD *vs* SD, between HFD groups and between SD groups, *p<0.05). Data are presented as mean ± standard error of the mean (SEM). Additional graphs show the comparison between male and female mice for each parameter by calculating the fold changes for each HFD mouse compared to the mean of their respective control group (HFD/SD) at time d28 and w14. Data were compared with unpaired *t*-tests (*\*p*<0.05) and are presented as mean ± standard error of the mean (SEM). (C) Graphs show the mean distribution of the number of Sholl intersections as a function of the distance from the astrocyte soma for SD mice (grey curves) and HFD mice (blue and pink curves for male and female mice, respectively) at d28 (N=3-4 cells/hemi-ARC from 6 mice/group) and at w14 (N=3-4 cells/hemi-ARC from 4 mice/group). Values were compared with 2-way ANOVA (HFD x distance) followed by Sidak’s multiple comparisons test (HFD *vs* SD, *\*p*<0.05).

Proinflammatory microglia have been demonstrated to release inflammatory cytokines and especially undergo deramification characterized by the retraction of their processes with loss of complexity (Butovsky and Weiner, 2018). Thus, we performed a Sholl analysis of individual 3D-reconstructed cells. To this end, spheres with a radius of 5 µm were superimposed starting at the center of the soma, and the number of process intersections that each sphere encountered was measured with IMARIS software. The microglia of HFD-fed mice displayed a significant lower complexity level (indicated by a significant reduced average number of total Sholl intersections) than those of SD-fed mice, but this difference occurred earlier for F-HFD i.e., at d28 (*p(HFD)*<0.05) compared to M-HFD which was no present at d28 but at w14 (*p(HFD)*<0.05, **Fig. 10C**). Conversely, the astrocytes of HFD mice displayed a significantly greater degree of complexity than those of SD-fed mice at d28 for both the M-HFD (*p(HFD)*<0.05) and F-HFD (*p(HFD)*<0.001) groups, but this difference persisted at w14 only for M-HFD (*p(HFD)*<0.05, **Fig. 11C**). Interestingly, the mean peak number of Sholl intersections was reached at 15 µm from the soma for both the SD-fed and HFD-fed mice at d28 and w14, and the mean peak number was significantly different between F-SD and F-HFD starting at 10 µm (p<0.05) to 20 µm from the soma at d28 (p=0.0518, **Fig**. **11C**).

## 4. Discussion

In these two experiments, we investigated the impact of obesity on the hypothalamic response by using a DIO model. Indeed, DIO models with high-fat diet (HFD) consumption are commonly used to induce obesity in rodents (de Moura E dias et al., 2021). Starting at 7 weeks of age, the first cohort of C57Bl/6J male and female mice received either a SD or a HFD with 60% kcal from fat for 14 weeks. First, the monitoring of body weight and body composition allowed us to confirm the robustness of our model. Indeed, compared to control mice, both M- and F-HFD-fed mice gained significantly more weight and exhibited significant changes in body composition with a greater increase in fat mass.

Regarding sex differences in the response to HFD, we first noted that F-HFD mice exhibited lower absolute weight gain and fat mass level values than male mice at the end of 14 weeks. However, when we calculated the final body weight gain to initial weight ratio, we noticed that the normalized increase was similar between male and female mice, suggesting that the HFD had similar effect on male *versus* female mice, at least for these weight gain criteria. Nevertheless, the weight gain to initial weight ratio over the course of the 14 weeks exposure to HFD appeared to be different between male and female animals, with M-HFD mice weight gain appearing faster than F-HFD mice. These results pushed us to investigate different HFD exposure duration to investigate the dynamics of weight gain in male and female mice.

Still, given the different composition of the HFD diet when compared to control SD diet, we wanted to assess the impact of a HFD at this time point on regulatory molecules of energy homeostasis and food intake. We hypothesized HFD could present strong satietogenic potential due to its high caloric density (5.24 kcal/g *vs* 3.34 kcal/g for SD). This was supported by weekly food consumption per cage from week 1 to week 12. Indeed, we reported a significant main effect of HFD: a lower food intake expressed in g in HFD mice compared to SD but a higher caloric intake, which was more pronounced in males compared to females (**Supplementary Fig. S3A, B**). It has been documented that a HFD induces neuronal adaptations that affect central neuropeptides involved in mediating satiety (Hamamah et al., 2023). For instance, studies have reported that dietary fats activate orexigenic neurons coexpressing neuropeptide Y (NPY)/Agouti-related protein (AgRP) while also attenuating anorexigenic neurons coexpressing proopiomelanocortin (POMC)/cocaine- and amphetamine-related transcript (CART), which contributes to hyperphagia and eventually leads to obesity (Hamamah et al., 2023). Interestingly, we found opposite results in HFD-fed mice, with a downregulation of mRNA transcripts encoding orexigenic neuropeptides and an upregulation of those encoding POMC but only in the F-HFD group. Furthermore, the increase in fat mass in HFD mice was also accompanied by glucose intolerance and higher insuline and leptin plasma concentrations compared to SD mice (Lefebvre et al., 2024). Our findings regarding food intake data support a satietogenic effect of a HFD and could reflect adaptative feedback mechanisms following HFD or suggest a potential leptin dependant effect in our model. Indeed, leptin directly influences the secretion patterns of NPY and AgRP peptides as well as POMC and CART peptides, either by inhibiting or activating neurons after binding to leptin receptors in the arcuate nucleus of the hypothalamus (Hamamah et al., 2023). Results in female mice could be explained by their greater leptin sensitivity compared to that of males (Chowen et al., 2018). Indeed, females are known to have higher oestrogen levels, and it was reported that it exerts anorexigenic effects by enhancing leptin sensitivity (Brown & Clegg, 2010; Clegg et al 2006). In particular, 17β-estradiol (E2) enhances POMC neurotransmission to inhibit NPY/AgRP neurons (Stincic et al 2018).

Growing evidence indicates that HFD consumption induces hypothalamic inflammation accompanied by reactive gliosis (Bhusal et al., 2022; Sewaybricker et al., 2022). Thus, we aimed to investigate the neuroinflammation profile in our model by performing quantitative real-time PCRs targeting several markers and using immunohistochemistry methods.

In contrast to previous studies (Bhusal et al., 2022), we did not observe significantly greater mRNA expression levels of proinflammatory cytokines in HFD-fed mice. However, we noted that mRNA transcript levels in the hypothalamus extracts were overall extremely low and often below the threshold of experimental detection (Ct>35) both for SD and HFD mice, thus representing a quantitative bias. Despite that fact, we found that the expression of mRNAs encoding CD11b and IBA1 increased and decreased, respectively, in the F-HFD group. These markers are expressed by both microglia and perivascular myeloid cells as well as infiltrating macrophages. Therefore, these data require careful interpretation. Indeed, nonspecific targeting does not allow us to identify the cell origin of these findings. We could exclude the possibility of potential infiltration of peripheral macrophages because they are not presumed to be present in the healthy CNS, unless an increase in blood‒brain barrier (BBB) permeability occurred in our model. To address this issue, we targeted the purinergic receptor P2RY12, which has been identified as a specific microglial marker allowing to distinguish microglia from macrophages (Reddaway et al., 2023; Toledano Furman et al., 2020). More precisely, this receptor, expressed on ramified processes, is involved in microglial membrane ruffling and chemotaxis (Toledano Furman et al., 2020). Thus, its overexpression contributes to process extensions followed by migration, the first step of neuroinflammation (Gómez Morillas et al., 2021). Nevertheless, the P2RY12 mRNA expression did not differ between the SD and HFD groups in both sexes, suggesting that the variations of CD11b and IBA1 are not related to microglia. However, the decrease in the level of the IBA1 mRNA in the F-HFD group was confirmed by immunofluorescence staining for the IBA1 protein in the ARC. These results could be associated with a particular microglial phenotype called “dark” microglia. Indeed, this population is particularly dominant in the hypothalamus and is known to downregulate IBA1 and P2RY12 but strongly expresses CD11b, which is involved in synaptic pruning (Stratoulias et al., 2019). Taken together, these findings allowed us to provide better precisions on the immunologically active profile of microglia in female mice after 14 weeks of HFD.

The deleterious effects of chronic HFD feeding on body changes and metabolic disturbances appeared to be uncoupled with the presence of neuroinflammation, at least from our RT‒qPCR studies, which must be confirmed by further analyses. We hypothesized that neuroinflammation could be more prominent at earlier stages of the exposure to HFD. To address this question, we performed a kinetic to analyse of the hypothalamic response during the initial phase of HFD feeding.

In this second cohort, male and female mice were exposed to HFD for 3, 7, 14 or 28 days. As suggested by the first study, male mice responded earlier to HFD exposure in terms of body weight gain and fat storage (starting from d3) compared to female mice where the response was significant only at d28. These differences were evident both in term of absolute weight gain and normalized weight when compared to initial weight, showing that HFD does indeed triggers sex-dependent responses in earlier stage of HFD exposure for these criteria. This delayed response and protection from short-term HFD consumption has already been highlighted in the literature (Huang et al., 2020). Indeed, this latter study reported that consumption of a HFD leads to hyperphagia in male rodents, while female mice have greater energy expenditure and are more resistant to body weight gain. In addition, it has been reported that females metabolic response allowed to maintain a normal body weight (Chowen et al., 2018).

Similar to the first cohort (w14), the consumption of a short-term HFD quickly led to a downregulation of NPY and AgRP (from d3), which was maintained at d28 for both sexes. Moreover, we found an increase in POMC expression at d3 and d14 for M- and F-HFD mice, respectively, which interestingly corresponded to the beginning of the greater body weight gain and fat mass for M-HFD than those of M-SD mice. This could be a potential compensatory mechanism to restore energy homeostasis and limit weight gain.

Taking into account that some of their data were obtained in rats (Thaler et al., 2012), our RT‒ qPCR analysis did not reveal a complex “on-off-on” pattern with significantly elevated hypothalamic levels of IL-6, IL-1β, TNF-α or glial markers in HFD mice as demonstrated in Thaler’s study. However, we observed a trend toward increased IL-6 (until d14), and CD11b (from d14) in M- and F-HFD mice, respectively. Furthermore, Fizz-1 levels were significantly decreased at d7 in M-HFD and at d3 in F-HFD, and the expression of Arg-1 tended to decrease in HFD mice. Taken together, these results only partially support the establishment of anti-inflammatory mechanisms in response to a HFD.

Hypertrophic and hyperplasic astrocytes are concurrently observed in the hypothalamus even 1 day after HFD feeding to reduce lipid overload (Thaler et al., 2012). Since GFAP is the hallmark intermediate filament, we expected to observe an increase in GFAP mRNA expression in HFD mice compared to that in SD groups. In contrast, we observed a downregulation during this kinetic experiment for both sexes. We could suppose a variability in expression is linked to the growth of mice or reflects a decrease in the size, number, and/or fibrous character of astrocytes. Immunostaining aimed to clarify these observations. In the same way that for the first study, we performed immunofluorescence staining within the ARC due to its proximity to the median eminence (ME), where the fenestrated vascular endothelium lacks a BBB. Indeed, among hypothalamic regions, the ARC provides a positional advantage to rapidly sense excessive fatty acids from the diet. In contrast with M-HFD and what was observed at w14, the number of IBA1+ cells seemed constant in female mice after a short-term HFD. In addition, we found a significant decrease in the number of GFAP+ cells at d3 in F-HFD. It must be taken into consideration that the counting was not carried out in the whole hypothalamus but within the ARC, which can explain these differences compared to RT‒qPCR data. Moreover, immunofluorescence assays have recently shown that the microglial signature varies according to hypothalamic nuclei and dietary fat content, particularly in mice (Mendes et al., 2021).

The classical markers, such as increased expression of IBA1 and GFAP or various cytokines, provide only indirect information about neuroinflammation and are insufficient to assess this process. Therefore, a detailed morphological analysis of glial cells was performed in order to provide additional and critical insights into hypothalamic adaptations under HFD. Using Sholl analysis, we reported significant differences between the SD-fed and HFD-fed mice. Regarding microglia, M-HFD mice showed a shift towards reduced complexity level compared to M-SD at w14, whereas this shift was already present at d28 for F-HFD. In contrast, Sholl analysis of astrocytes revealed greater complexity in HFD-fed mice than in SD-fed mice with anew earlier response to F-HFD. In addition, we reported an increase in the filament volume and the number and length of branches in the M-HFD and F-HFD groups at d28, respectively. Astrocytes along with microglia are highly dynamic cells and display a complex spectrum of morphotypes depending on the biological context as well as the local environment (Paolicelli et al., 2022). The more pronounced effect of HFD on astrocytes than on microglia is consistent with their involvement in several metabolic processes (glucose homeostasis maintenance, insulin and leptin sensing, etc.) (García-Cáceres et al., 2016). Indeed, astrocytes are the main site of fatty acid metabolism and oxidation in the brain (Chowen et al., 2018). Taken together with the mRNA expression analysis, these changes in astrocytic morphology (and supposedly activity) might be interpreted as metabolic adaptations caused by the introduction of excess lipids rather than an inflammatory response. Still, we observed a reduction in the complexity of microglial ramification which translate to a reactive phenotype. HFD has been showed to trigger neuroinflammation and to induce a morphological transition from a highly ramified to a deramified state, retracting their fine processes while simultaneously undergoing somatic hypertrophy (Reddaway et al., 2023). In this last experiment, we reported differences in glial parameters under HFD conditions in a time and sex dependent manner, but it remains unclear whether these changes translate to proinflammatory and/or neuroprotective mechanisms or altered glial function. This study demonstrates that a strong inflammatory hypothalamic profile is not required to induce obesity in mice and questions the role of the modulation of glial cell function in the induction of HFD phenotype. Therefore, further investigation is needed to better characterize this relationship. This could be achieved using a multidimensional integration of transcriptomic, metabolomic, proteomic and epigenetic approaches.

Because mice were co-housed (4 animals per cage), we were not able to measure neither locomotor activity nor energy expenditure for each animal. However, CRH mRNA levels only decreased in the M-HFD group, which is known to increase energy expenditure (Luquet, 2008). Similar studies assessing these parameters have shown divergent findings. Indeed, some data suggest that females are more active than males but in rats (Eckel & Moore, 2004). Studies carried out in C57Bl/6 mice reported that male mice reduced physical activity in response to a HFD with unaffected food intake, while female mice showed increased food intake without changes in physical activity and exhibited increased energy expenditure. Some studies did not show sex differences in locomotor activity, which is also not affected by HFD (Maric et al, 2022). This variability between studies can be linked to inconsistencies in study designs using a variety of diet types, fat content with different ω6/ω3 ratios (Sanchez et al., 2024), onset of intervention and duration of diet, making elucidating what variables cause the observed divergences in the literature challenging. Despite these limitations, our study emphasizes the crucial role of carefully examining sex differences and timing in DIO models.

## 5. Conclusion

In summary, the present study provides nuanced data regarding hypothalamic inflammation under HFD conditions. However, we support evident sexual dimorphism in response to short- and long-term HFD consumption, suggesting that the prevention and treatment of obesity should be considered in a sex-dependent manner.

## 6. Acknowledgements

We thank the PRIMACEN platform (HeRaCleS Inserm US51, CNRS UAR 2026, Université de Rouen Normandie, France) to the microscopic facilities.

## 8. Author’s contribution

VD, AG, PD, MC and LL, conceptualization; VD, CL, CE-B, MC and LL, formal analysis; VD, CL, CE-B, CS, CBF, CG and LL, investigation; VD, MC and LL original draft preparation; all authors writing review and editing.

## 9. Data availability

The data sets used and analyzed from the current study are available from the corresponding author upon reasonable request.

## Abbreviations

AgRP: Agouti-related protein
ARC: Arcuate nucleus
BDNF: Brain-derived neurotrophic factor
CRH: Corticotropin-releasing hormone
DIO: Diet-induced obesity
GFAP: Glial fibrillary acidic protein
HFD: High-fat diet
Iba1: Ionized calcium-binding adapter molecule 1
IL: Interleukin
iNOS: Inducible nitric oxide synthase
MC4R: Melanocortin-4 receptor
NPY: Neuropeptide Y
PBS: Phosphate-buffered saline
PFA: Paraformaldehyde
POMC: Proopiomelanocortine
P2RY12: Purinergic receptor
P2Y12 ROI: Region of interest
RT-qPCR: Reverse transcriptase quantitative polymerase chain reaction
SD: Standard diet
TNF: Tumor necrosis factor

## Supplementary Information

**Supplementary Table S1: Composition of the high fat diet.**

**Supplementary Table S2: Primer sequences used for RT-qPCR.**

**Supplementary Fig. S1: Visualization of glial cells by immunofluorescence in ARC after 14 weeks of SD or HFD.**

These representative images were taken within the ARC and show GFAP and IBA1 immunopositive cells in male (top panel) and female mice (bottom panel) after 14 weeks of SD or HFD feeding. Scale bar: 50 µm. 3V: third ventricle.

**Supplementary Fig. S2: Comparisons of body composition changes between male and female mice after the short-term HFD feeding.**

(A, B) Ratio of each mouse HFD / mean of respective SD group of fat mass (in g and % of body weight) and (C, D) lean mass (in g and % of body weight) measured by EchoMRI in C57Bl/6J male and female mice fed either SD or HFD feeding for up 28 days (experiment 2, n=4 SD for d3, d7, d14; n=12 SD for d28 and n=12 HFD/time). Values were compared with 2-way ANOVA (sex x time) followed by Sidak’s multiple comparisons test (\**p*<0.05, \*\**p*<0.01, \*\*\**p*<0.001, \*\*\*\**p*<0.0001). Data are presented as mean ± standard error of the mean (SEM).

**Supplementary Fig. S3: Weekly monitoring of food consumption in male and female mice over 12 weeks of diet.**

Average estimation of food consumed in grams and kilocalories (g and kcals) per day and per animal fed either SD or HFD in male mice (A) and female mice (B). N=6 cages/group, total of grams and kcals measured per cage were divided by the number of mice per cage (n=4/cage). Average values per cage were compared with 2-way ANOVA (HFD x time) followed by Sidak’s multiple comparisons test (\**p*<0.05, \*\**p*<0.01, \*\*\**p*<0.001, \*\*\*\**p*<0.0001). Data are presented as mean ± standard error of the mean (SEM). Male and female mice fed HFD per day ingested in average 3g (15.73 kcals) and 2.4g (12.71 kcals) compared to male and female mice fed with SD which per day consumed in average 4.1g (13.69 kcals) and 3.6g (12.16 kcals), respectively.

