## Supplementary table 1 for "Sex-dependent effects of a high-fat diet on the hypothalamic response in mouse"

**Supplementary Table S1: Composition of the high fat diet.**

| Formula #D12492 | gm % | kcal % |
| --- | --- | --- |
| Protein  Carbohydrate  Fat  Total  kcal/gm | 26.2  26.3  34.9  5.24 | 20  20  60  100 |
| Ingredients | **gm** | **kcal** |
| Casein, 30 Mesh  L-Cystine  Corn Starch  Maltodextrin  Sucrose  Cellulose, BW200  Soybean Oil  Lard  Mineral Mix S10026  DiCalcium Phosphate  Calcium Carbonate  Potassium Citrate, 1 H2O  Vitamin Mix V10001  Choline Bitartrate  FD&C Blue Dye #1  Total | 200  3  0  125  68.8  50  25  245  10  13  5.5  16.5  10  2  0.05  773.85 | 800  12  0  500  275.2  0  225  2205  0  0  0  0  40  0  0  4057 |
| Fatty acid profile | **gm** | **%** |
| Saturated  Monounsaturated  Polyunsaturated | 81.7  91.2  81.0 | 32.2  35.9  31.9 |
| n6  n3  ratio | 73.7  5.9  13.9 |  |
