## Supplementary table 2 for "Sex-dependent effects of a high-fat diet on the hypothalamic response in mouse"

**Supplementary Table S2: Primers used for real time PCR.**

|  | Forward primer (5’🡪3’) | Reverse primer (5’🡪3’) |
| --- | --- | --- |
| *Gapdh* | CATCACTGCCACCCAGAAGA | AAGTCGCAGGAGACAACfiCT |
| *Actb* | CTCTGGCTCCTAGCACCATGAAG | GTAAAACGCAGCTCAGTAACAGTCCG |
| *Il1b* | CCCAAAAGATGAAGGGCTGC | AAGGTCCACGGGAAAGACAC |
| *Il6* | CACTTCACAAGTCGGAGGCT | CTGCAAGTGCATCATCGTTGT |
| *Tnfa* | TGTCTACTCCTCAGAGCCCC | TGAGTCCTTGATGGTGGTGC |
| *Iba1* | ATCAACAAGCAATTCCTCGATGA | CAGCATTCGCTTCAAGGACAT |
| *Gfap* | TGGAACAGCAAAACAAGGCG | CTGTCTATACGCAGCCAGGT |
| *P2ry12* | ACGGACACTTTCCCGTATCC | AAGTTCCCAAAGCCCTCTGT |
| *Npy* | CTGCGACACTACATCAATCT | CTTCAAGCCTTGTTCTGG |
| *Pomc* | CCTCCTGCTTCAGACCTCCA | GGCTGTTCATCTCCGTTGC |
| *Agrp* | ACTGAAGGGCATCAGAAGGC | TTGAAGAAGCGGCAGTAGCA |
| *Cd11b* | ATGGACGCTGATGGCAATACC | TCCCCATTCACGTCTCCCA |
| *Fizz1* | CCTGAGATTCTGCCCCAGGAT | TTCACTGGGACCATCAGCTGG |
| *Inos* | CCGAAGCAAACATCACATTCA | GGTCTAAAGGCTCCGGGCT |
| *Arg1* | GTTCCCAGATGTACCAGGATTC | CGATGTCTTTGGCAGATATGC |
| *Mc4r* | TCTCTATGTCCACATGTTCCTG | GGGGCCCAGCAGACAACAAAG |
| *Crh* | TAAAGAAAATGTGGCCCCAAGG | CTTCCACTGCAGCTCCAAATAA |
| *Bdnf* | TGTGACAGTATTAGCGAGTGGGT | TACGATTGGGTAGTTCGGCATT |
