## Supplementary figures and images for "Sex-dependent effects of a high-fat diet on the hypothalamic response in mouse"

### Supplementary figure 1

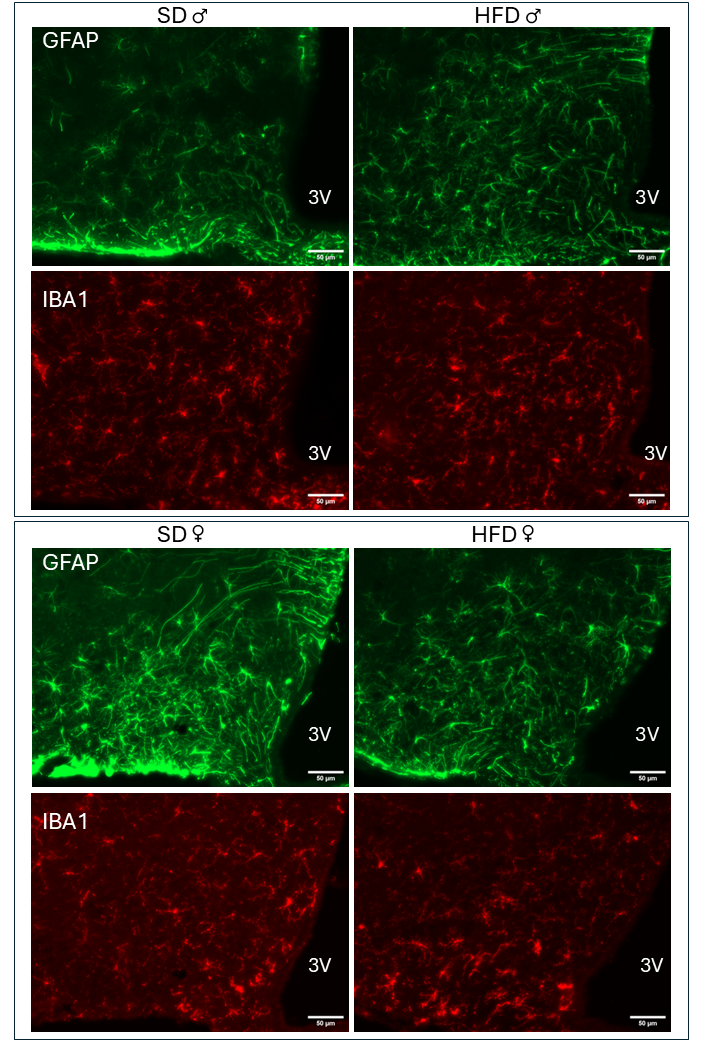

### Supplementary figure 2

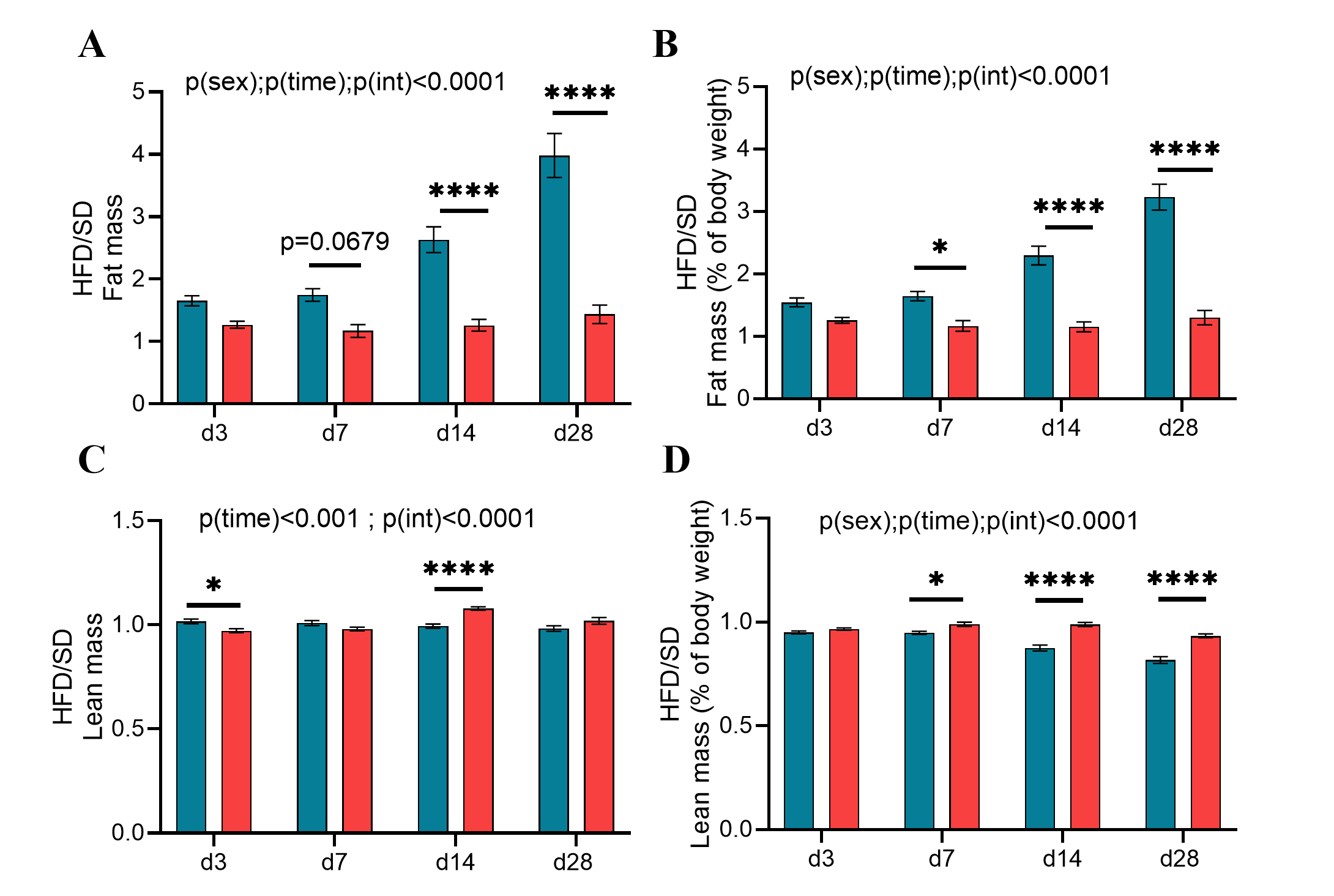

### Supplementary figure 3

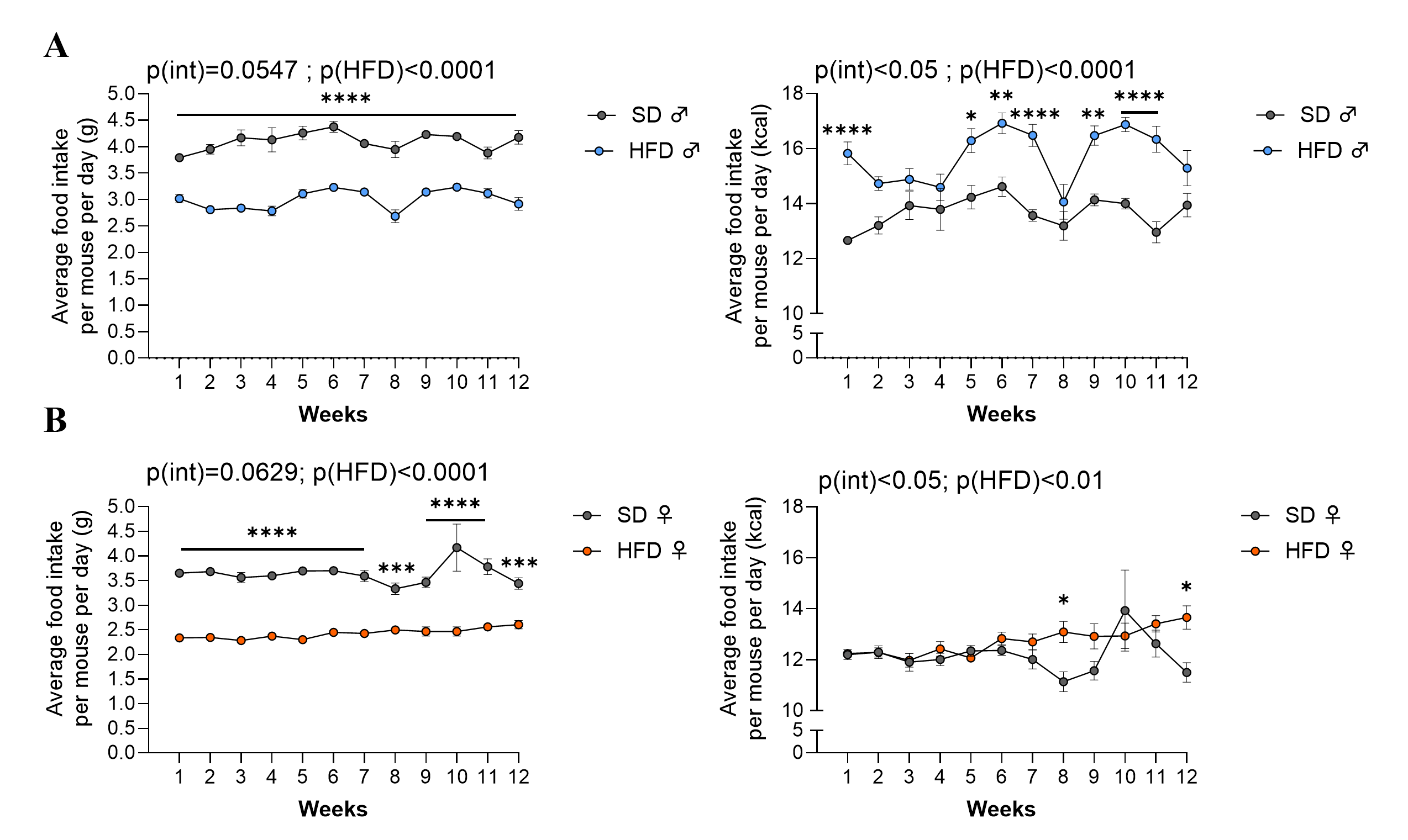
